# An intestinally secreted host factor promotes microsporidia invasion of *C. elegans*

**DOI:** 10.1101/2021.07.12.452088

**Authors:** Hala Tamim El Jarkass, Calvin Mok, Michael R. Schertzberg, Andrew G. Fraser, Emily R. Troemel, Aaron W. Reinke

**Affiliations:** Department of Molecular Genetics, University of Toronto, Toronto, ON, Canada; Division of Biological Sciences, University of California, San Diego, La Jolla, California, United States of America; Donnelly Centre, University of Toronto, Toronto, ON, Canada

## Abstract

Microsporidia are ubiquitous obligate intracellular pathogens of animals. These parasites often infect hosts through an oral route, but little is known about the function of host intestinal proteins that facilitate microsporidia invasion. To identify such factors necessary for infection by *Nematocida parisii*, a natural microsporidian pathogen of *Caenorhabditis elegans*, we performed a forward genetic screen to identify mutant animals that have a Fitness Advantage with *Nematocida* (Fawn). We isolated four *fawn* mutants that are resistant to *Nematocida* infection and contain mutations in *T14E8.4*, which we renamed *aaim-1* (Antibacterial and Aids invasion by Microsporidia). Expression of AAIM-1 in the intestine of *aaim-1* animals restores *N. parisii* infectivity and this rescue of infectivity is dependent upon AAIM-1 secretion. *N. parisii* spores in *aaim-1* animals are improperly oriented in the intestinal lumen, leading to reduced levels of parasite invasion. Conversely, *aaim-1* mutants display both increased colonization and susceptibility to the bacterial pathogen *Pseudomonas aeruginosa* and overexpression of *AAIM-1* reduces *P. aeruginosa* colonization. Competitive fitness assays show that *aaim-1* mutants are favoured in the presence of *N. parisii* but disadvantaged on *P. aeruginosa* compared to wild type animals. Together, this work demonstrates how microsporidia exploits a secreted protein to promote host invasion. Our results also suggest evolutionary trade-offs may exist to optimizing host defense against multiple classes of pathogens.

## Introduction

Microsporidia are a large group of obligate intracellular parasites that infect most types of animals.^1^ These ubiquitous parasites possess the smallest known eukaryotic genoms, and are extremely reliant on their host as a result of the loss of many genes involved in metabolism and energy production.^2, 3^ Microsporidia can have a large impact on the evolution of their hosts, as infection with microsporidia often leads to a reduction in host offspring and the effect of this selective pressure has resulted in resistant animals within a population.^4, 5^ Microsporidia are currently a major threat to many commercially important species such as honeybees and shrimp.^6, 7^ Many species also infect humans and infections in immunocompromised individuals can result in lethality.^8^ Despite their ubiquitous nature, effective treatment strategies are currently lacking for these poorly understood parasites.^9^

Microsporidia infection begins with invasion of host cells. They possess fascinating invasion machinery, a unique structure known as the polar tube.^10^ This apparatus, resembling a long thread, is often coiled within a dormant spore. However, once inside of a host, and in proximity to the tissue of interest, the polar tube rapidly emerges or “fires”, releasing the infectious material (the sporoplasm) which is deposited intracellularly either through direct injection, or through the internalization of the sporoplasm.^10, 11^

A number of microsporidia proteins have been demonstrated to play important roles during invasion by insect- and human-infecting species of microsporidia.^10^ For example, spore wall proteins can interact with host cells through the recognition of sulfated glycosaminoglycans, heparin binding motifs, integrins, and proteins on the cell surface. ^12–17^ In *Encephalitozoon* species, polar tube proteins (PTP) can mediate interactions with the host. For instance, O-linked mannosylation on PTP1 has been demonstrated to bind mannose binding receptors, whereas PTP4 interacts with the transferrin receptor (Trf1).^11, 18–20^ Additionally, the sporoplasm surface protein, EhSSP1, binds to an unknown receptor on the cell surface.^21^ These proteins on the spore, polar tube, and sporoplasm have all been shown to promote microsporidia adhesion or invasion of host cells in culture systems, but the role of these proteins during animal infection is unclear.

The nematode *Caenorhabditis elegans* is infected in its natural habitat by several species of microsporidia, and frequently by *Nematocida parisii.*^22–24^ This species infects the intestinal cells of *C. elegans,* which possess similarity to those of mammalian cells, making this animal both a relevant tissue and model to study these infections in vivo.^24, 25^ Infection of *C. elegans* by *N. parisii* begins when spores are consumed by the worm, where they then pass through the pharynx into the intestinal lumen and fire, depositing sporoplasms inside of intestinal cells. Within 72 hours the sporoplasm will divide into meronts, which differentiate into spores, that then exit the animal, completing the parasite’s life cycle.^26, 27^ Infection with *N. parisii* leads to reduced fecundity and premature mortality in *C. elegans*.^24, 26^ Several mutants have been shown to affect proliferation and spore exit.^28, 29^ Immunity that can either prevent infection or clear the pathogen once infected has also been described.^4, 27, 30–32^ In contrast, very little is known about how *N. parisii* invades *C. elegans* intestinal cells. Almost all of the microsporidia proteins known to facilitate invasion of host cells are not conserved in *N. parisii* and although host invasion factors described in other species are present in *C. elegans*, there is no evidence that they are being used by microsporidia during invasion of *C. elegans.*^11^

To understand how microsporidia invade animal cells, we performed a forward genetic screen to identify host factors that promote infection. We identified a novel, nematode-specific protein, AAIM-1, whose loss of function confers resistance to microsporidia infection. This protein is expressed in intestinal cells, secreted into the intestinal lumen, and is necessary to ensure proper spore orientation during intestinal cell invasion. In addition, we show that AAIM-1 limits bacterial colonization of pathogenic *Pseudomonas aeruginosa*. Strikingly, AAIM-1 plays opposing roles on host fitness in the face of pathogenesis. The utilization of a host factor critical for bacterial defense reflects a clever strategy to ensuring microsporidia’s reproductive success.

## Results

### A forward genetic screen identifies *aaim-1* as being necessary for *N. parisii* infection

To identify host factors needed for infection by microsporidia, we carried out a forward genetic screen using a *C. elegans* model of *N. parisii* infection. We took advantage of the previously described phenotypes of *C. elegans* displaying reduced fitness when infected with *N. parisii,* including reduced progeny production and stunted development.^26, 27, 33, 34^ We mutagenized animals and subjected their F2 progeny to *N. parisii* infection. After infecting populations for five subsequent generations, we selected individual worms containing embryos, indicating increased fitness in the presence of infection (see Methods). We identified four independent isolates that when exposed to *N. parisii* reproducibly had higher fractions of animals containing embryos compared to wild type (N2). We named these isolates Fitness Advantage With *Nematocida* (*fawn* 1-4) (Figure S1a).

As *C. elegans* that are less infected with *N. parisii* produce more progeny, we hypothesised that these *fawn* mutants would be resistant to *N. parisii* infection^27^. To determine this, we grew the three isolates with the strongest phenotype, *fawn* 1-3, in the presence and absence of *N. parisii*, and stained each population of worms with the chitin binding dye, Direct-yellow 96 (DY96), at 72 hours post infection (hpi). DY96 allows for the visualization of chitinous microsporidia spores as well as worm embryos (Figure 1a). In the absence of infection, there is no difference in the fraction of *fawn-2* and *fawn-3* animals developing into adults containing embryos (gravid adults), although *fawn-1* has a modest defect. In comparison, all three *fawn* isolates generate significantly more gravid adults than N2 animals in the presence of infection (Figure 1b). We next examined the fraction of animals in each strain containing intracellular microsporidia spores and observed that all three *fawn* isolates display significantly fewer numbers of spore-containing worms (Figure 1c).

**Figure 1:**
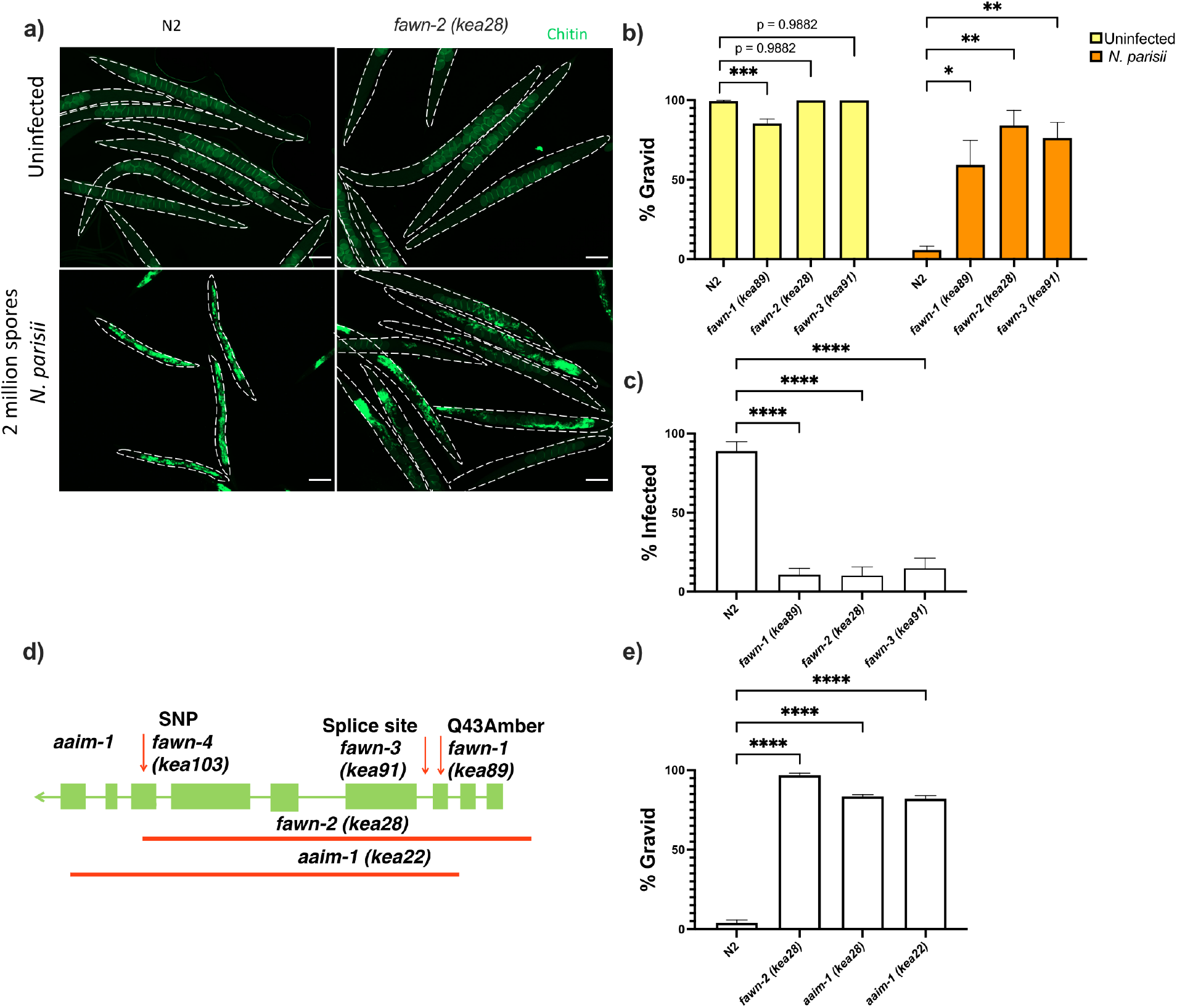
Mutations in *aaim-1* result in resistance to *N. parisii* infection. (a-c, and e) L1 stage wild-type (N2) and *aaim-1* mutant animals were infected with either a high dose (a, b, and e) or a very lose dose (c) of *N. parisii*, fixed at 72 hours, and stained with direct- yellow 96 (DY96). (a) Representative images stained with DY96, which stains *C. elegans* embryos and microsporidia spores. Scale bars, 100 μm. (b and e) Graph displays percentage of gravid worms. (c) Percentage of worms that contain newly formed *N. parisii* spores. (d) Schematic depicting the nature and location of the different *aaim-1* alleles. Boxes represent exons, and connecting lines represent introns. Arrows depict point mutations, and the solid red lines depict deletions. *fawn-2 (kea28)* has a 2.2 kb deletion and *aaim-1 (kea22)* has a 2.3 kb deletion. *fawn-1 (kea89)* carries a C127T, Q43Stop mutation, *fawn-3 (kea91)* carries a G221A splice site mutation and *fawn-4 (kea103)* carries a C1286T, A429V mutation in *aaim-1*. (b,c, and e) Data is from three independent replicates of at least 90 worms each. Mean ± SEM represented by horizontal bars. P- values determined via one-way ANOVA with post hoc. Significance defined as: * p < 0.05, ** p < 0.01, *** p < 0.001, **** p < 0.0001.

### These results suggest that *fawn* mutants are missing an important factor for efficient microsporidia infection

To identify the causal mutations underlying the Fawn phenotype, we used a combination of whole-genome sequencing and genetic mapping. We generated F2 recombinants and performed two rounds of infection with microsporidia, selecting for gravid animals. After each round we used molecular inversion probes to determine the region of the genome linked to the causal mutation.^35^ This revealed strong signatures of selection on the left arm of chromosome X in all three *fawn* isolates and absent in N2 (Figure S1b). Analysis of whole genome sequencing showed that all four *fawn* isolates contained different alleles of *T14E8.4*, which we named *aaim-1* (Antibacterial and Aids Invasion by Microsporidia-1) for reasons described below (Figure 1d). We validated the role of *aaim-1* in resistance to infection using several additional alleles: an independent allele *aaim-1* (*ok295*), carrying a large gene deletion in both *aaim-1* and *dop-3,* and a CRISPR-Cas9 derived allele, *aaim-1 (kea22),* that contains a large gene deletion. Both of these alleles displayed a fitness advantage when infected with *N. parisii* (Figure 1d,e, S1c,d). These data demonstrate that *aaim-1* is the causative gene underlying the *fawn* 1-4 infection phenotypes. In subsequent experiments we utilized both *aaim-1 (kea22)*, and *fawn-3 (kea28),* carrying a 2.2 kb deletion in *aaim-1,* which was outcrossed to N2 six times (hereafter referred to as *aaim-1 (kea28)*).

### *aaim-1* is expressed in the pharynx and intestine, and secretion is important for function

AAIM-1 is a poorly characterized protein that does not possess any known or conserved domains. Homologs of the protein exist in both free-living and parasitic nematodes (Figure S2). To further characterize the role of AAIM-1 during *N. parisii* infection, we generated transgenic extrachromosomal lines of *C. elegans* carrying a reporter transgene of GFP under control of the *aaim-1* promoter. GFP fluorescence was observed in the terminal bulb of the pharynx as well as the posterior of the intestine throughout development (Figure 2a). Embryos and L1 animals display additional expression in the arcade cells of the pharynx (Figure 2a, S3a).

**Figure 2:**
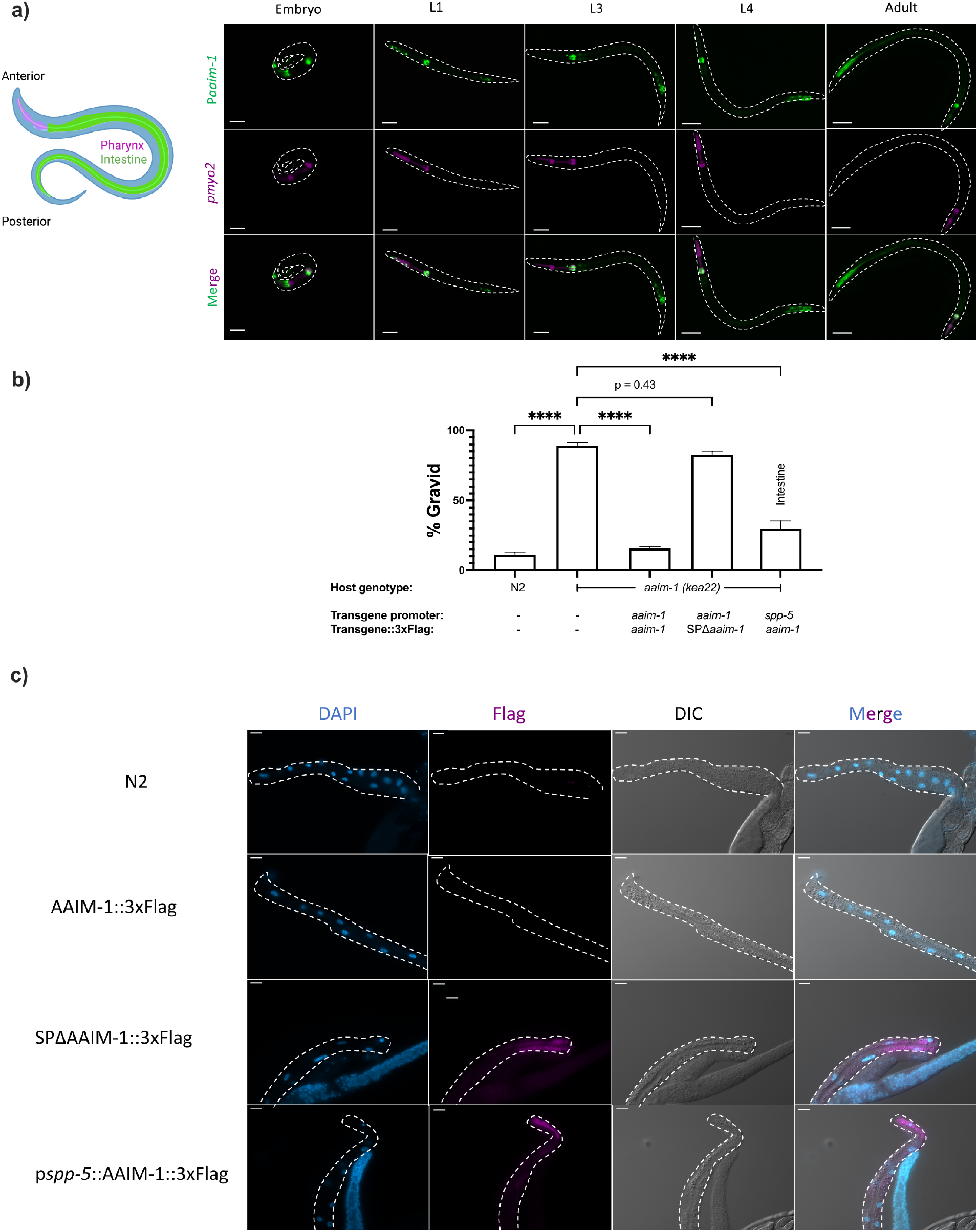
AAIM-1 is secreted from intestinal cells. (a) Wild-type worms containing an extrachromosomal array expressing GFP from the *aaim-1* promoter and mCherry (labelled in magenta) in the pharyngeal muscles were imaged at the embryo, L1, L3, L4, and adult stage. Embryo, L1, and L3 animals were imaged at 40x, scale bar 20 μm and L4 and adult animals were imaged at 20x, scale bar 50 μm. L1 to L4 animals are oriented anterior to posterior and the adult animal is oriented posterior to anterior from left to right. Schematic made with Biorender.com (b) N2, *aaim-1, and aaim-1* expressing extrachromosomal arrays were infected with a medium-2 dose of *N. parisii*. Graph displays percentage of gravid worms. Data is from three independent replicates of at least 90 worms each. Mean ± SEM represented by horizontal bars. P-values determined via one-way ANOVA with post hoc. Significance defined as **** p < 0.0001 (c) Intestines (denoted by dashed lines) of 72-hour post- L1 adults were dissected and stained using anti-Flag (magenta) and DAPI (blue). Images taken at 40x, scale bar 20 μm.

The first 17 amino acids of AAIM-1 are predicted to encode a signal peptide.^36^ This suggests that AAIM-1 may be secreted into the pharyngeal and intestinal lumen, the extracellular space through which *N. parisii* spores pass before invading intestinal cells. To test which tissues AAIM-1 functions in and if secretion is important for function, we generated a series of transgenic worms expressing extrachromosomal arrays (Key resources table). First, we generated transgenic *aaim-1 (kea22)* animals expressing AAIM-1 tagged on the C-terminus with a 3x Flag epitope. Transgenic animals expressing AAIM-1 under its native promoter complement the ability of *aaim-1 (kea22)* animals to develop into adults in the presence of a high amount of *N. parisii* spores (Figure 2b). A construct expressing GFP or GFP::3xFlag does not influence this phenotype nor does the presence of the epitope tag impair the ability of AAIM-1 to rescue the mutant phenotype (Figure S3b). We next generated a signal peptide mutant allele of AAIM-1 missing the first 17 amino acids *(SPΔaaim-1),* which is unable to complement the *aaim-1 N. parisii* infection phenotype. In contrast, AAIM-1 expressed from an intestinal-specific promoter (*spp-5)*^37^ can rescue the infection phenotype of *aaim-1 (kea22)* (Figure 2b).

To determine where AAIM-1 localizes, we dissected the intestines from transgenic worms and performed immunofluorescence using anti-Flag antibodies. We were unable to detect expression of AAIM-1::3xFlag when expressed from its endogenous promoter. However, we observed protein expression in the intestinal cells of animals expressing AAIM-1::3xFlag from a strong, intestinal specific-promoter or when the signal peptide was removed (Figure 2c). We did not observe AAIM- 1::3xFlag localized in the extracellular space of the intestinal lumen, possibly due to rapid turnover of intestinal contents or due to loss from dissection of the intestines.^38^ The increased expression in the signal peptide mutant suggests an accumulation of protein that is unable to be secreted. Taken together, these data demonstrate that AAIM-1 is secreted and acts within the intestinal lumen to promote *N. parisii* infection.

### AAIM-1 is only necessary for microsporidia infection at the earliest larval stage

*N. parisii* infection of *C. elegans* can occur throughout development, but several forms of immunity towards microsporidia have been shown to be developmentally regulated.^4, 27^ To determine if *aaim-1* mutant animals display developmentally restricted resistance to infection, we infected *fawn* 1-3 at the L1 and L3 stage. For these experiments we took advantage of another intestinal-infecting species of microsporidia, *Nematocida ausubeli*, which has a more severe effect on *C. elegans* fecundity, allowing us to determine fitness defects after the L1 stage.^4, 23, 26^ *fawn* isolates are resistant to *N. ausubeli* as seen by an increase in the fraction of gravid adults in the population after exposure to a medium dose of *N. ausubeli* (Figure 3a). When we initiated infections at the L3 stage of growth, *fawn* isolates do not have increased resistance, and instead exhibit wild-type levels of susceptibility (Figure 3b). To rule out the possibility that this L1 restricted phenotype was the result of exposure to sodium hypochlorite treatment, which we used to synchronize worms, we exposed embryos that were naturally laid by adults within a two-hour window to *N. parisii* infection. Animals synchronized in this manner still display a robust resistance to *N. parisii* (Figure S4c). Thus, resistance to infection in *aaim-1* mutants is developmentally restricted and AAIM-1 is utilized by several species of microsporidia.

**Figure 3:**
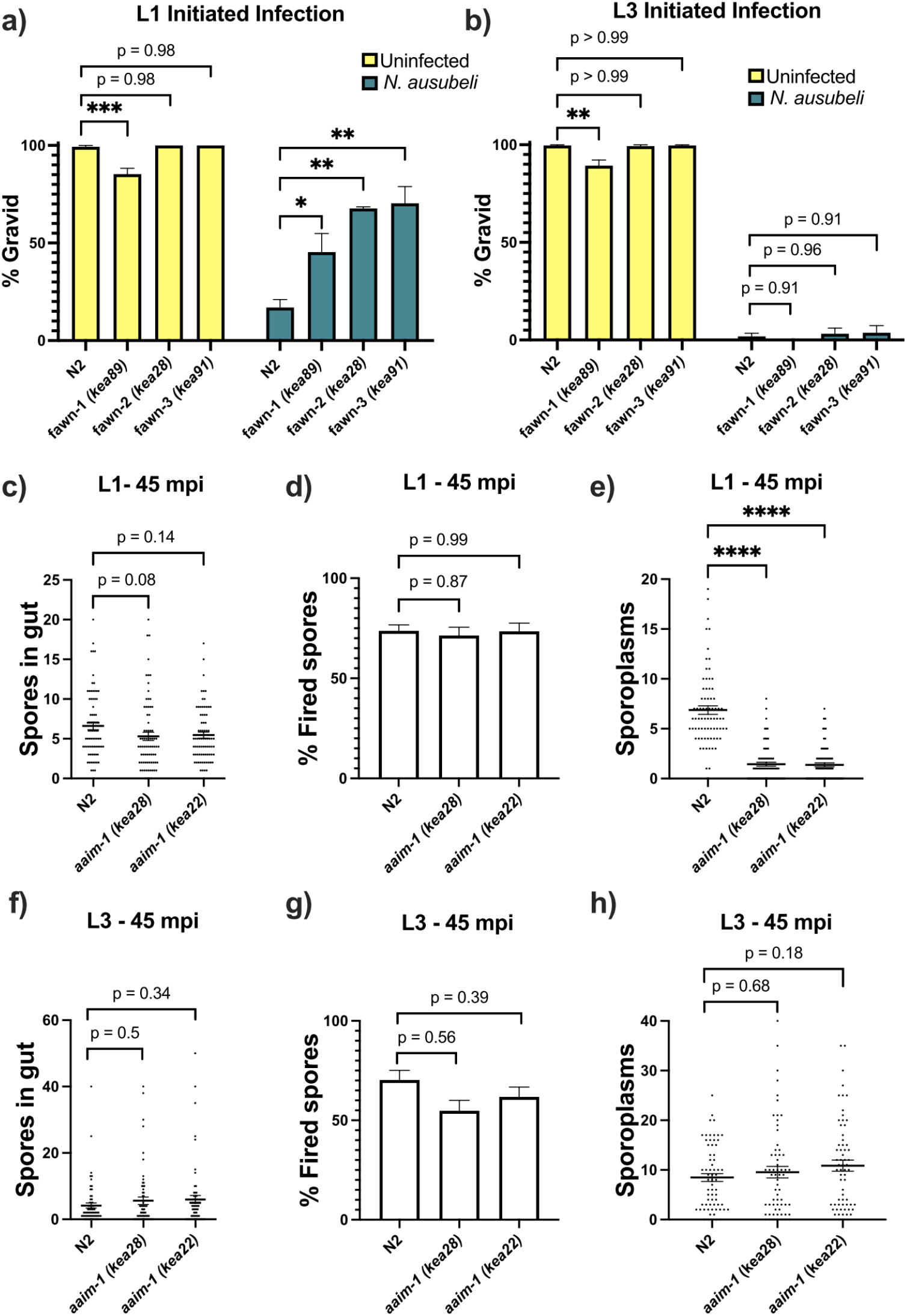
*aaim-1* mutants are resistant to microsporidia at the earliest larval stage due to spore misfiring. **(a-b)** N2 and *aaim-1* mutants were infected with a medium dose of *N. ausubeli* at either the L1 stage for 72 hours (a) or a high dose of *N. ausubeli* at the L3 stage for 48 hours (b) Graph displays percentage of gravid worms. (c-f) N2 and *aaim-1* animals were infected with a medium-3 dose of *N. parisii* for 45 minutes at L1 (c-e) or L3 (f-h), fixed, and then stained with DY96 and an *N. parisii* 18S RNA fish probe. The number of spores per animal (c,f), the percentage of spores fired (d,g), and the number of sporoplasm per worm (e,h) are displayed. Data is from three independent replicates of at least 100 worms each (a-b) or 20-30 worms each (c-h). (a-h) Mean ± SEM represented by horizontal bars. P-values determined via one-way ANOVA with post hoc. Significance defined as: * p < 0.05, ** p < 0.01, *** p < 0.001, **** p < 0.0001.

### AAIM-1 is needed for efficient invasion of intestinal cells

Resistance to infection could be the result of a block in invasion, proliferation, or through the destruction of the parasite. To test the mechanism of resistance in *aaim-1* mutants, we performed pulse-chase infection assays at the L1 and L3 stage of development.^4, 27^ Here, we treated animals with a medium-1 dose (as defined in Table S1) of *N. parisii* for 3 hours, washed away any un- ingested spores, and then replated the animals in the absence of spores for an additional 21 hours. We then used an 18S RNA Fluorescent In Situ Hybridization (FISH) probe to detect *N. parisii* sporoplasms, which is the earliest stage of microsporidia invasion. In our *fawn* 1-3 isolates we detect less invasion at 3hpi compared to N2 (Figure S4a). However, there was no reduction in the number of infected animals between 3hpi and 21 hpi, indicating that pathogen clearance was not occurring. This defect in invasion was not present at the L3 stage, providing further support that resistance is restricted to the L1 stage in *aaim-1* mutants (Figure S4b). A reduction in invasion could be due to a feeding defect, leading to a reduction in spore consumption. To test rates of consumption, we measured the intestinal accumulation of fluorescent beads. We find that *aaim-1* alleles displayed wild-type levels of bead accumulation, unlike the feeding defective strain *eat-2 (ad465)* (Figure S4d).

For *N. parisii* to invade host cells, spores must first enter the intestinal lumen and fire their polar tube.^27^ To test if *aaim-1* mutants have defects in spore entry or spore firing, we infected animals for either 45 minutes or 3 hours, at the L1 and L3 stages. We then fixed and stained animals with both an *N. parisii* 18S RNA FISH probe and DY96 and quantified the number of spores present in the intestinal lumen of animals. Here, *aaim-1* animals infected for 45 minutes or 3 hours at L1 or L3 contained similar amounts of spores as N2 animals (Figure 3c,f, S5a,d). The percentage of fired spores present within these animals is also not significantly different at either developmental stage (Figure 3d,g, S5b,e). We then counted the number of sporoplasms per animal and observed significantly fewer invasion events in *aaim-1* mutant animals infected at L1 (Figure 3e, S5c). In contrast, the number of sporoplasms in L3 stage *aaim-1* alleles are similar to that observed in the N2 strain (Figure 3h, S5f). These results demonstrate that the *N. parisii* invasion defect in *aaim-1* mutants is not caused by differences in spore firing or accumulation. Instead, these results suggest that spores are misfiring, leading to unsuccessful parasite invasion.

### AAIM-1 plays a role in promoting proper spore orientation

To determine how AAIM-1 promotes *N. parisii* invasion, we further examined the invasion process. We pre-stained spores with Calcofluor white (CFW) and assessed their orientation relative to the intestinal apical membrane using the apical membrane marker PGP-1::GFP in L1 worms infected for 45 minutes (Figure 4a). In N2 animals, 32.4% of spores are angled relative to the apical membrane. In contrast, spores in an *aaim-1* mutant were angled 14.3% of the time (Figure 4b). Several host factors that promote microsporidia invasion cause adherence to host cells.^11^ To determine if AAIM-1 influences the location of spores relative to intestinal cells in *aaim-1* mutants, we measured the perpendicular distance from the center of a parallel spore to the apical membrane of the intestine. Surprisingly, parallel spores in *aaim-1* alleles were significantly closer to the apical membrane (0.29 μm) than those in N2 (0.34 μm) (Figure 4c). In agreement with resistance being developmentally restricted, *aaim-1* mutants display wild-type spore orientations and distances from the membrane when infections were initiated at the L3 stage (Figure 4d,e). The width of the intestinal lumen at the L1 stage does not differ significantly between N2 and *aaim-1* mutants, however, L3 animals generally possess wider intestinal lumens (Figure S5g,h). Thus, taken together these results suggest that AAIM-1 plays a distinct role in the intestinal lumen at L1 to promote proper spore orientation, through maintaining an appropriate distance and angle to the apical membrane, resulting in successful invasion.

**Figure 4:**
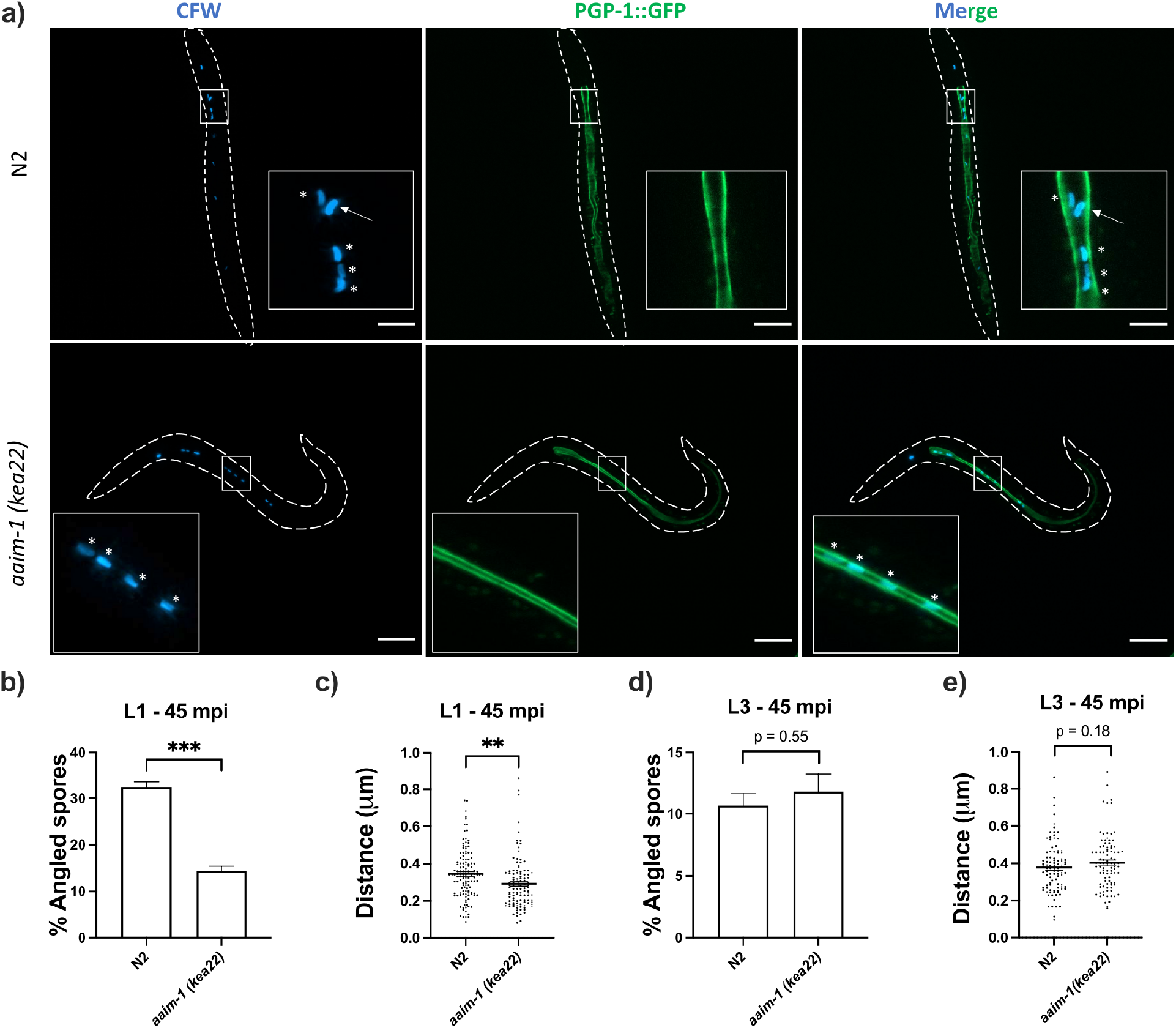
Spores in *aaim-1* mutants display improper orientation and distance to the apical intestinal membrane. (a-e) PGP-1::GFP and *aaim-1(kea22);* PGP-1::GFP animals were infected with a very high dose of Calcofluor white (CFW) pre-stained *N. parisii* spores for 45 minutes at either the L1 stage (a-c) or the L3 stage (d-e). (a) Representative images of live animals containing stained spores (blue) relative to the apical intestinal membrane (GFP). Arrow indicates an example of an angled spore and asterisks indicate parallel spores. Images taken at 63x, scale bar 20 μm. (b, d) Percentage of angled spores. Data is from three independent replicates of at least 90 spores each. (c, e) Distance from the center of each spore to the intestinal apical membrane. Data is from three independent replicates of at least 25 spores each. Mean ± SEM represented by horizontal bars. P-values determined via unpaired Student’s t-test. Significance defined as ** p < 0.01, *** p < 0.001.

### AAIM-1 inhibits intestinal colonization by *Pseudomonas aeruginosa*

Interestingly, *aaim-1* has been shown to be upregulated by a variety of different fungal and bacterial pathogens, including *P. aeruginosa*. ^39, 40^ Using our transcriptional reporter strain, we sought to confirm this and determine if microsporidia infection could also induce *aaim-1* transcription. N2 animals carrying a transcriptional reporter (p*aaim-1*::GFP::3xFlag) were exposed to *N. parisii, P. aeruginosa* PA14, or *E. coli* OP50-1, and the levels of GFP quantified when grown on these pathogens for 72 hours from the L1 stage, or for 24 hours from the L4 stage. Infection by either *N. parisii* or *P. aeruginosa* PA14 resulted in the upregulation of *aaim-1* as detected by an increase in the GFP signal (Figure 5a, S6f).

**Figure 5:**
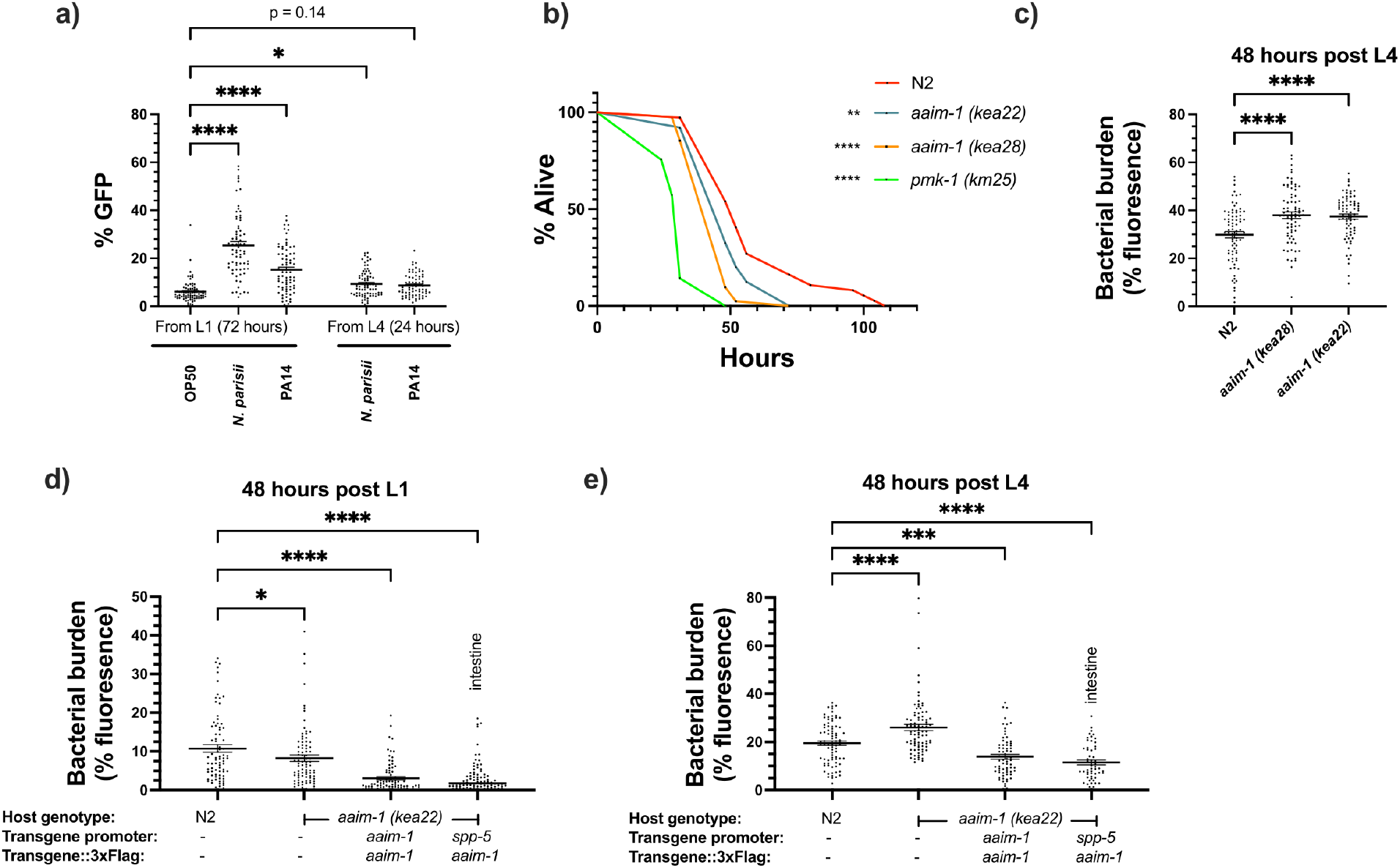
*aaim-1* is upregulated by *N. parisii and P. auerginosa* and *aaim-1* animals are susceptible to infection by *P. aeruginosa*. (a) Expression of p*aaim-*1::GFP::3xFlag in response to infection with either PA14 or *N. parisii* for either 72 hours from L1 or 24 hours from L4. Data is from three independent replicates of at least 18-25 worms each. Every point represents a single worm. Percentage GFP was measured as the percentage of the animal containing GFP via FIJI. (b) L4 stage N2, *aaim-1,* and *pmk-1 (km25)* animals were plated on full lawns of *P. aeruginosa* PA14::DsRed and the percentage of animals alive was counted over the course of 96 hours. TD_50_ : N2 48 hours, aaim-1 *(kea28)* 44 hours, *aaim- 1 (kea22)* 33 hours, and *pmk-1 (km25)* 28 hours. Three independent replicates were carried out, and a representative replicate is displayed. At least 37 worms were quantified per strain. P-values determined via Log-rank (Mantel-Cox) test. Significance defined as * p < 0.05, ** p < 0.01. (c-e) N2, *aaim-1,* or *aaim-1* animals with different extrachromosomal arrays were plated on PA14::DsRed at either the L1 stage (d) or L4 stage (c,e) for 48 hours. Bacterial burden was measured as the percentage of the animal containing PA14::DsRed. Data is from three independent replicates of 20-30 worms each. Every point represents a single worm. Mean ± SEM represented by horizontal bars. P- values determined via two-way (a) or one-way ANOVA(c-e) with post hoc. Significance defined as * p < 0.05, ** p < 0.01, *** p < 0.001, **** p < 0.0001.

Previously, an *aaim-1* deletion strain, RB563 *(ok295),* was shown to display reduced survival on lawns of *P. aeruginosa* PA14.^41^ The enhanced susceptibility previously reported was attributed to *dop-3,* which is also partially deleted in RB563 *(ok295)*. ^41^ To determine if *aaim-1* mutants are susceptible to pathogenic bacterial infection, we assayed the survival of L4 stage worms in *P. aeruginosa* PA14 slow killing assays. A mutant in the p38 MAPK pathway (*pmk-1*) was used as control for susceptibility to PA14^42^. We observed reduced survival in *aaim-1* alleles, although not to the same extent as the *pmk-1* mutant (Figure 5b, S6a,b,c). In contrast to significant susceptibility to the Gram-negative *P. aeruginosa, aaim-1* mutants do not display enhanced susceptibility to the Gram-positive bacterium *Staphylococcus aureus* NCTC8325, suggesting specificity of AAIM-1 to PA14 infection (Figure S7a).

Lethality in slow killing assays is a result of *P. aeruginosa* accumulation within the intestinal lumen.^43, 44^ To investigate if *aaim-1* alleles displayed higher levels of bacterial burden, animals were grown on lawns of PA14::DsRed at the L1 or L4 stage for 48 hours. *aaim-1* mutant alleles exposed as L4s, but not L1s, displayed higher bacterial burden relative to N2 (Figure 5c, S6d,e). To test if intestinal expression of *aaim-1* was sufficient to limit bacterial colonization, transgenic *aaim-1 (kea22)* overexpressing AAIM-1::3xFlag from the endogenous or an intestinal-specific promoter were exposed to lawns of PA14::DsRed. When grown for 48 hours at the L1 or L4 stage, bacterial burden was significantly reduced, relative to N2 (Figure 5d, e). These results indicate that AAIM-1 plays a role in limiting bacterial colonization, and its loss results in reduced survival due to hyper-colonization of the intestinal lumen.

### Fitness of *aaim-1* animals is dependent upon microbial environment

To investigate how *aaim-1* alleles can influence population structure, we set up competitive fitness assays. A *C. elegans* strain with a fluorescent marker (RFP::ZNFX-1) was co-plated with N2 or *aaim-1* mutants on *E. coli* OP50-1, *N. parisii* or *P. aeruginosa* PA14. Animals were grown for 8 days, such that the population was composed of adult F1s and developing F2s. On *E. coli* OP50- 1, there is equal representation of N2 and *aaim-1* mutants in the population (Figure 6a). This is consistent with *aaim-1* mutants not having a developmental delay (Figure 1b) or a decrease in longevity (Figure S7b). In contrast, growth on *N. parisii* resulted in *aaim-1* alleles outcompeting the N2 strain. Conversely, *aaim-1* mutants on *P. aeruginosa* PA14 did significantly worse, being underrepresented in the population compared to N2 (Figure 6a). Interestingly, wild isolates of *C. elegans* do not carry any obvious loss of function alleles of *aaim-1* suggesting that natural conditions have selected for its retention (Figure S8). ^45^

**Figure 6:**
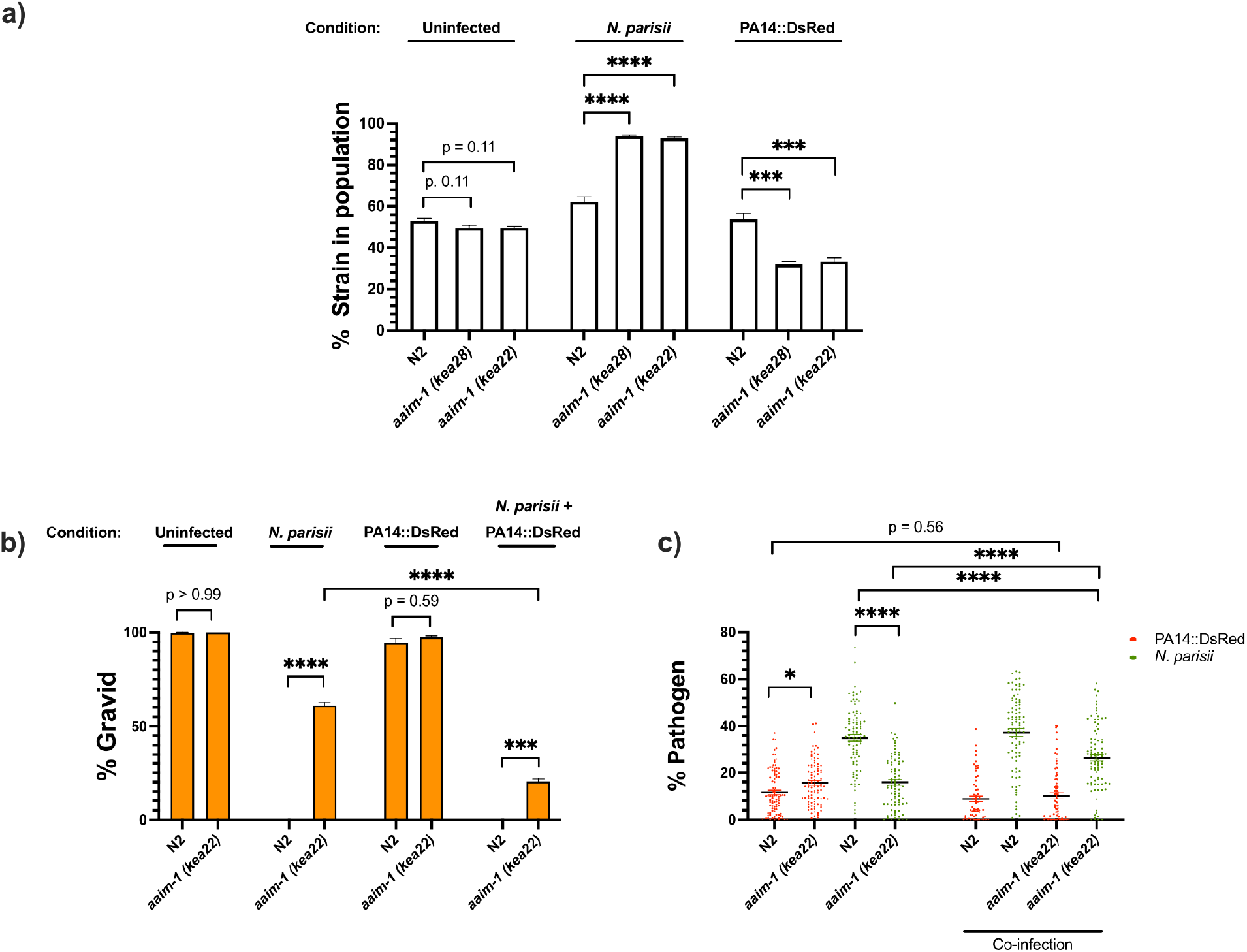
*aaim-1* alleles display enhanced fitness on *N. parisii*, but reduced fitness on *P. aeruginosa*. (a) Competitive fitness assays performed with a fluorescently marked strain (RFP::ZNFX1) mixed with either N2 or *aaim-1* mutants. These mixed populations of animals were plated at the L1 stage on either *E. coli*, a medium-2 dose of *N. parisii*, or on *P. aeruginosa*. After 8 days, the fraction of animals that did not display fluorescent germ granules was counted. Data is from three independent replicates of 20-270 worms each. (b,c) L1 stage N2 and *aaim-1* animals were either uninfected or infected with a maximal dose of *N. parisii*. These infected and uninfected population of animals were then washed and placed on either *E. coli* or PA14::DsRed. After 69 hours, animals were fixed and stained with DY96. Data is from three independent replicates of

Given the opposing fates of *aaim-1* mutants on *N. parisii* and *P. aeruginosa,* we investigated the effects of co-infection. Animals were infected with a maximal dose of *N. parisii* for 3 hours, prior to placement on lawns of PA14. For infections with a single pathogen, we observed similar results as before whereby *aaim-1* mutants have increased fitness in the presence of *N. parisii* and display lower levels of parasite burden but have increased bacterial accumulation when grown on PA14. In the presence of both pathogens, populations of *aaim-1* mutants display fewer gravid adults and increased amounts of *N. parisii* spores. (Figure 6b,c). These results suggests that coinfection with *N. parisii* and *P. aeruginosa* has synergistically negative effects on the fitness of *C. elegans*.

## Discussion

To identify host factors needed for microsporidia infection, we isolated mutants from a forward genetic screen that have a fitness advantage when challenged with *N. parisii* infection. This screen identified mutants in the poorly understood protein AAIM-1 (previously T14E8.4). Here, we demonstrate that this protein both promotes microsporidia invasion and limits colonization by pathogenic bacteria. Although we were unable to visualize the localization of secreted AAIM-1, our genetic and infection experiments strongly suggest that this protein acts in the intestinal lumen where both microsporidia invasion and bacterial colonization take place. The key role that AAIM-1 plays in immunity is further exemplified by its transcriptional regulation in response to infection (Figure 7).

**Figure 7:**
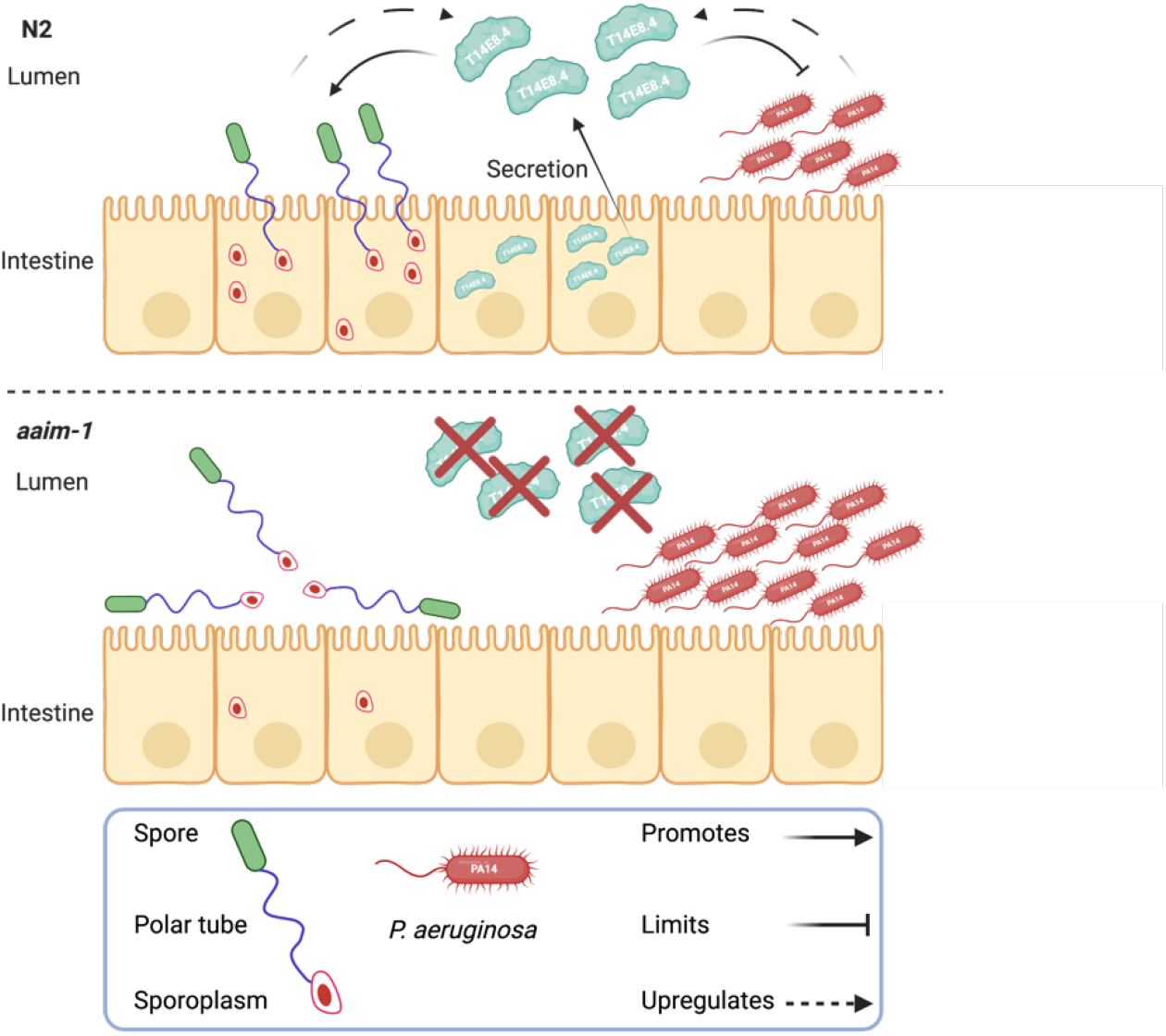
Secreted AAIM-1 functions in the intestinal lumen to limit bacterial colonization but is exploited by microsporidia to ensure successful invasion of intestinal cells. AAIM-1 is secreted from intestinal cells, where the protein limits bacterial colonization in the lumen. Additionally, AAIM-1 is parasitized by *N. parisii* spores to ensuring successful orientation and firing during intestinal cell invasion. Infection by either of these two pathogens results in the upregulation of AAIM-1. Figure made with Biorender.com.

The processes by which microsporidia invade host cells are poorly understood. We show that *N. parisii* spores are often angled in wild-type *C. elegans*, suggesting that successful invasion requires a particular spore orientation. In the absence of AAIM-1, spores are more often parallel to the intestinal lumen, where spores may fire without the successful deposition of the sporoplasm inside an intestinal cell. In contrast to previously described host and microsporidia proteins involved in invasion, AAIM-1 does not appear to be involved in promoting adhesion to the surface of host cells.^10, 11^ Instead, AAIM-1 ensures an adequate distance of spores from the intestinal membrane, possibly allowing spores to be able to properly orient themselves to ensure successful host cell invasion. *N. parisii* spores are ∼2.2 μm long by ∼0.8 μm wide and the average width of the intestinal lumen at the L1 stage is ∼0.6 μm.^23^ Therefore, at the L1 stage spores may not be able to move freely, but at the L3 stage, where AAIM-1 is not needed for invasion, there is less of a constraint on spore movement as the luminal width increases to ∼1.3 μm. Alternatively, the developmentally restricted role of AAIM-1 could be due stage-specific expression of other factors that work along with AAIM-1 to promote microsporidia invasion. Together, our results highlight the power of studying microsporidia invasion in the context of a whole animal model.

Several lines of evidence suggest that AAIM-1 plays a role in protecting animals against *P. aeruginosa*. First, *aaim-1* is upregulated in the intestine in response to PA14 exposure. Second, overexpression of AAIM-1 significantly decreases PA14 burden in the intestine. Third, loss of *aaim-1* leads to enhanced susceptibility and increased PA14 colonization. Fourth, competition assays show reduced reproductive fitness of *aaim-1* mutants on PA14. The survival phenotype of *aaim-1* mutants (∼78% survival compared to wild type) is modest compared to loss of the p38 MAPK pathway (∼51% survival compared to wild type). However, in competitive fitness assays, *aaim-1* mutants are ∼60% less represented than wild type in the F2 generation. Taken together, our data suggests that in addition to promoting microsporidia invasion, AAIM-1, at least in part, limits bacterial colonization and decreases susceptibility to *P. aeruginosa*.

*C. elegans* employs a variety of proteins to protect against bacterial infection. Many of these proteins belong to several classes of antimicrobial effectors used to eliminate and prevent colonization by pathogenic bacteria^46^, are upregulated upon infection, and predicted to be secreted.^47, 48^ One class of secreted proteins that is known to have immune functions and prevent bacterial adherence are the mucins. These large, glycosylated secreted proteins are upregulated during *C. elegans* infection and their knockdown alters susceptibility to *P. aeruginosa* infection. ^49, 50^ AAIM-1 has many predicted mucin-like O-glycosylation sites on serine and threonine residues.^51–53^ Thus, one possibility is that AAIM-1 may be functionally analogous to mucins, preventing the adhesion of microbes to the surface of intestinal cells. As AAIM-1 does not contain any known or conserved domains, further work will be necessary to determine its exact biochemical function.

*C. elegans* lives in a microbially dense environment containing a wide variety pathogens that *C. elegans* has evolved immunity towards.^23, 54–57^ Although loss of *aaim-1* provides a fitness advantage to *C. elegans* when grown in the presence of microsporidia, obvious loss of function alleles are not present in wild isolates sequenced thus far. Additionally, *aaim-1* mutants do not have observable defects when grown on non-pathogenic *E. coli*. This is in contrast to mutations in *pals-22* or *lin-35*, which negatively regulate the transcriptional response to infection and provide resistance to microsporidia infection when mutated, but at the cost of reduced reproductive fitness^27, 58^. Loss of *aaim-1* disadvantages *C. elegans* when grown on *P. aeruginosa*, but not *S. aureus,* suggesting AAIM-1 does not broadly promote resistance to all bacterial pathogens. These findings demonstrate that there is a trade-off in host defense between microsporidia and some pathogenic bacteria. The opposing functions of *aaim-1* with different pathogens adds to the limited set of known examples of trade-offs that constrain the evolution of host defense to multiple biotic threats^59, 60^.

## Methods

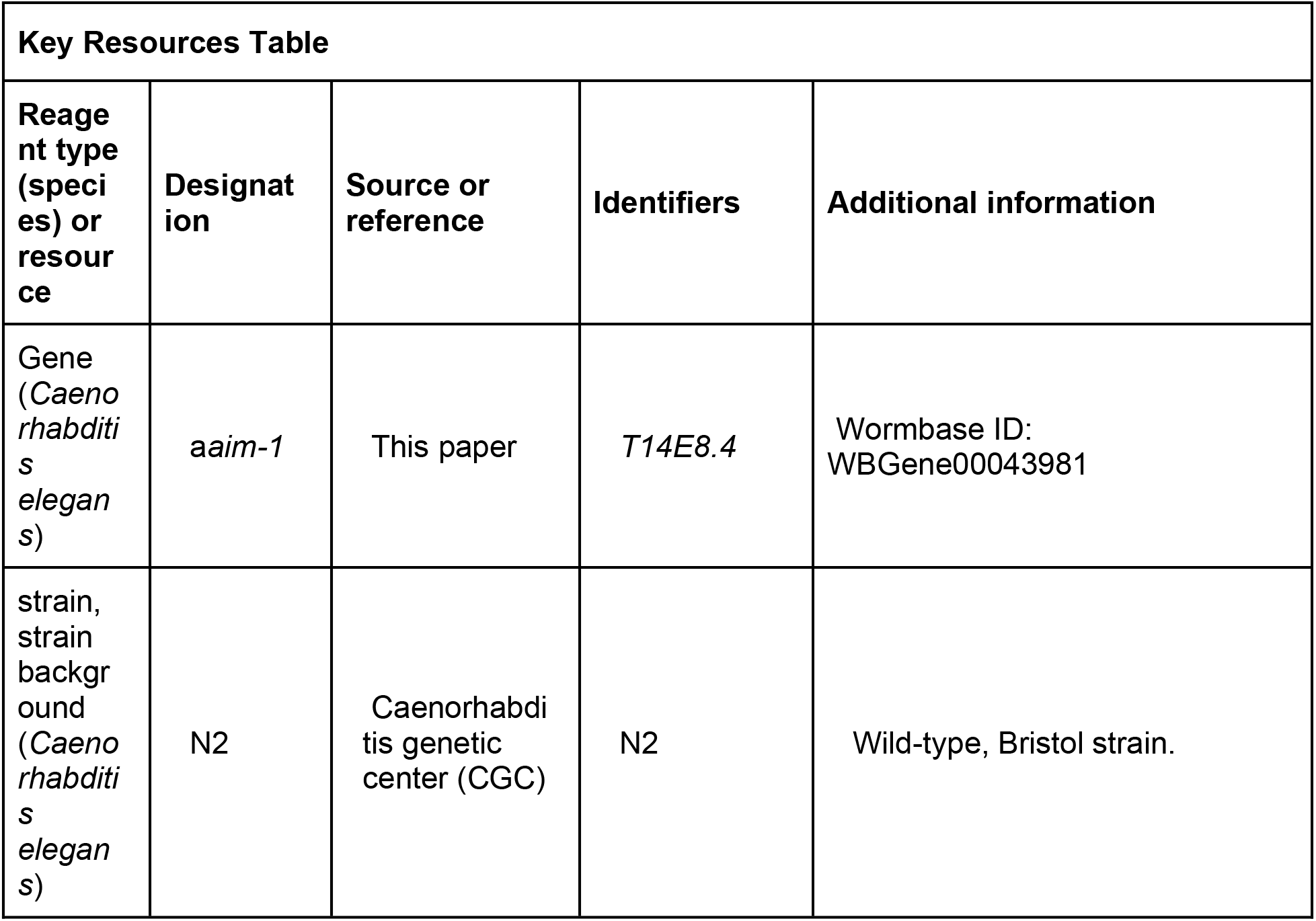
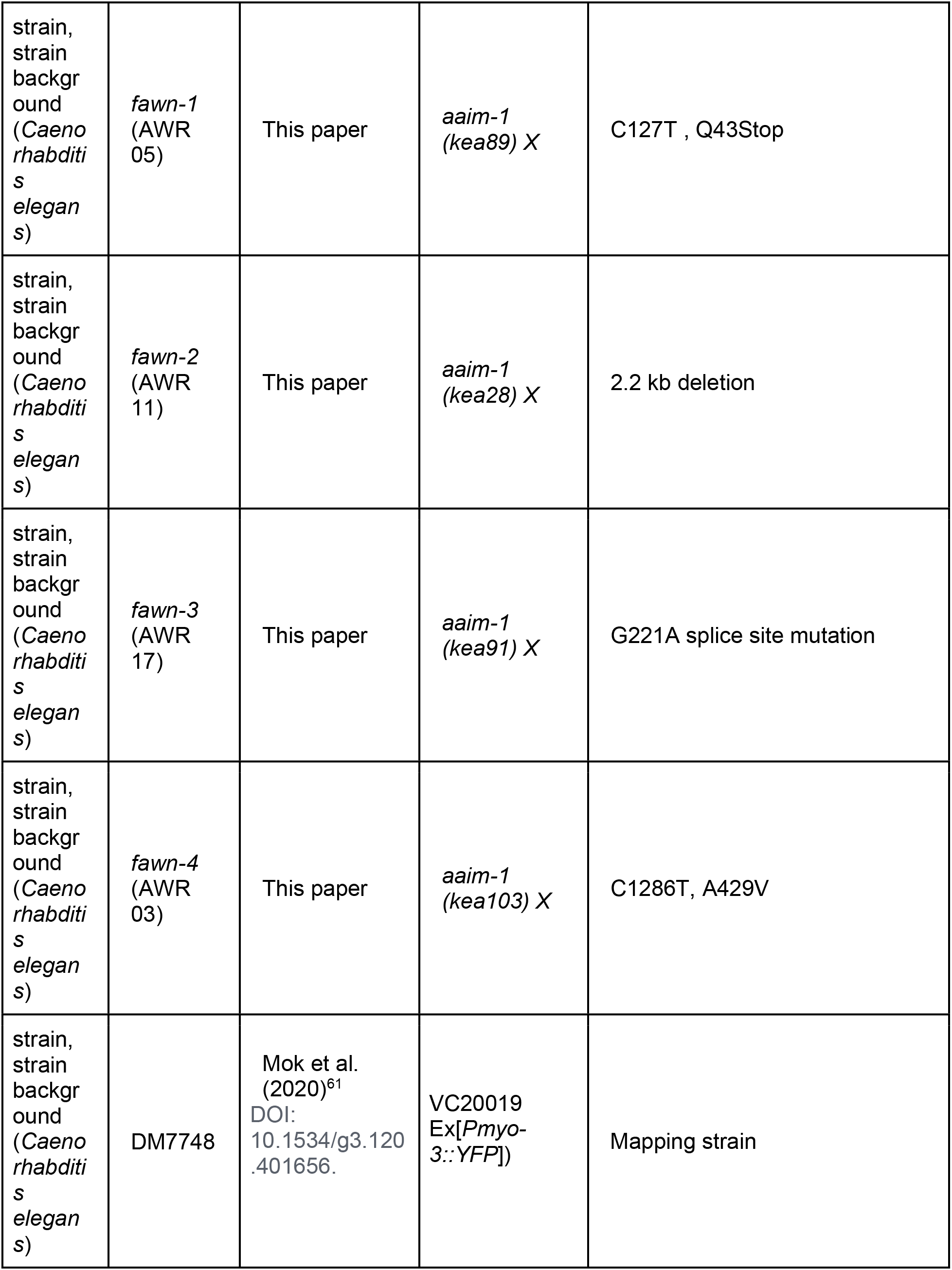
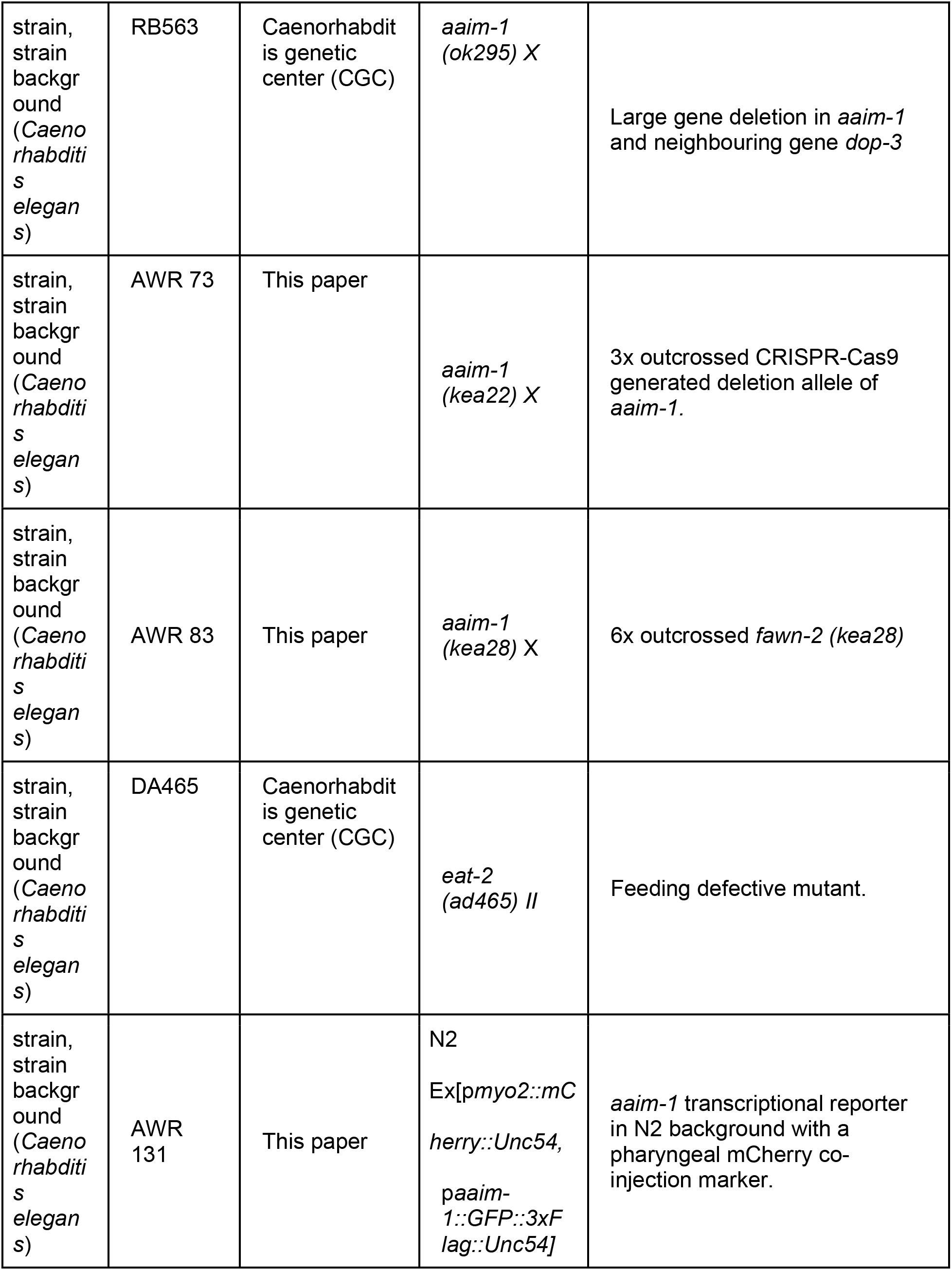
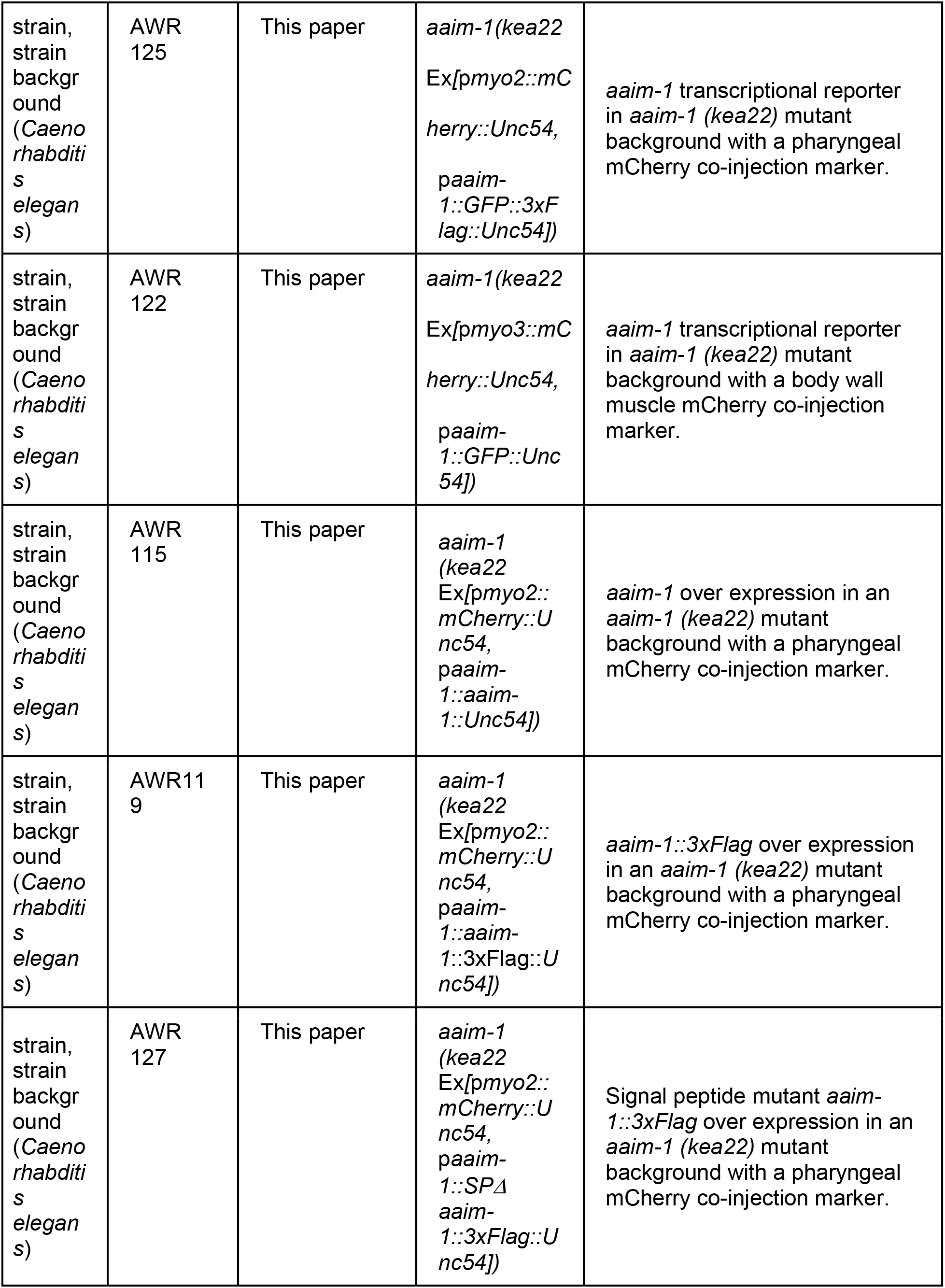
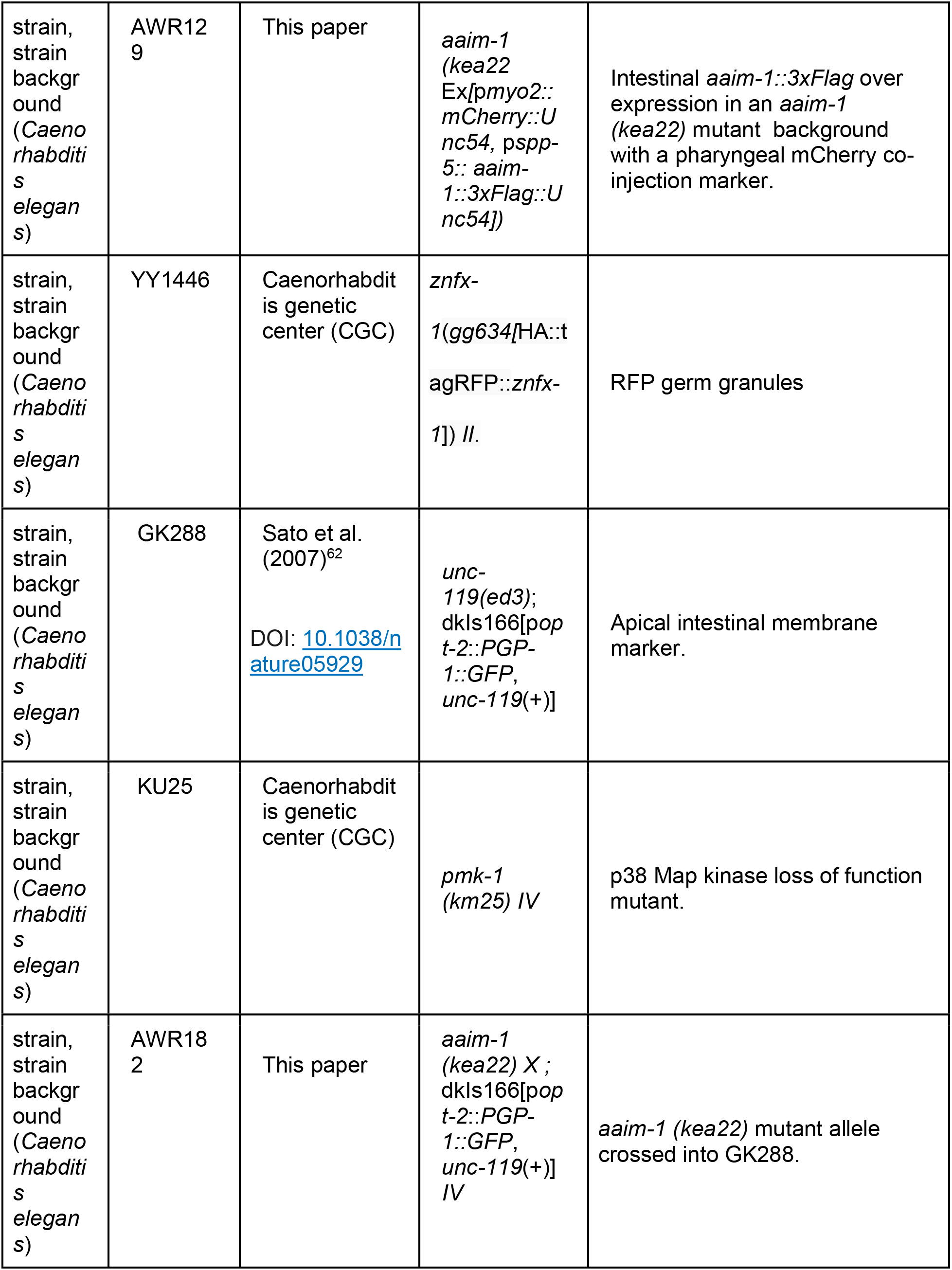
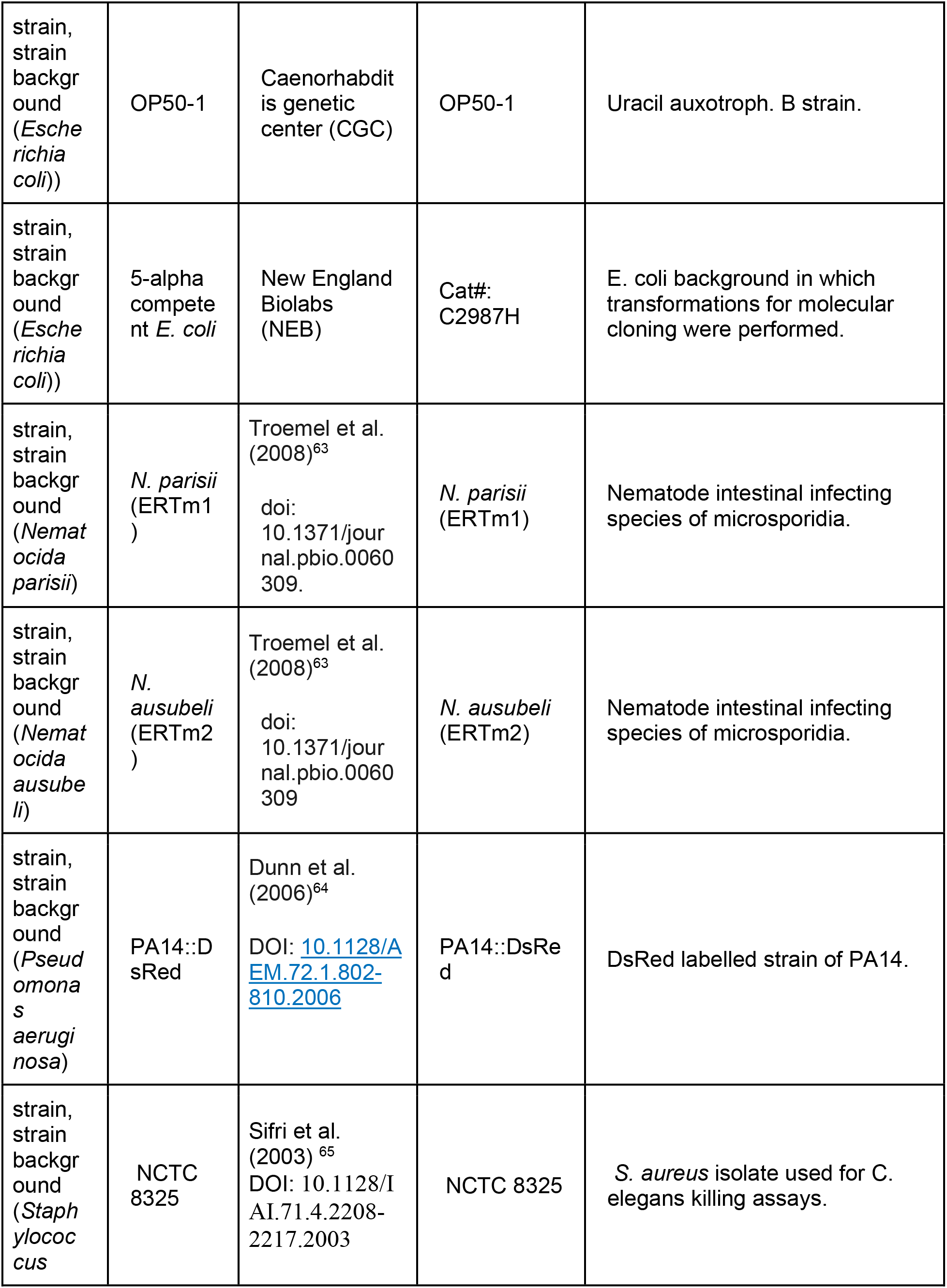
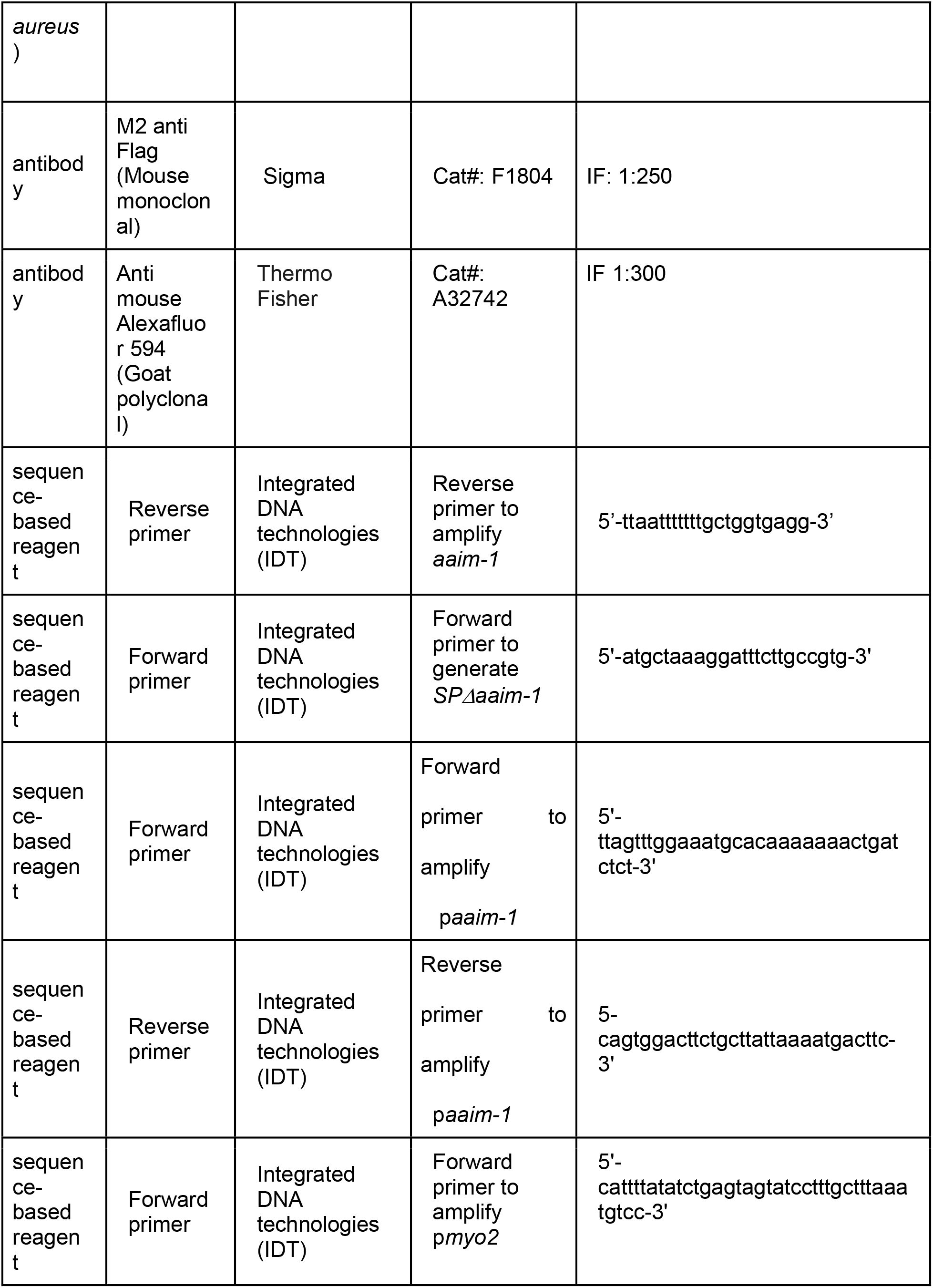
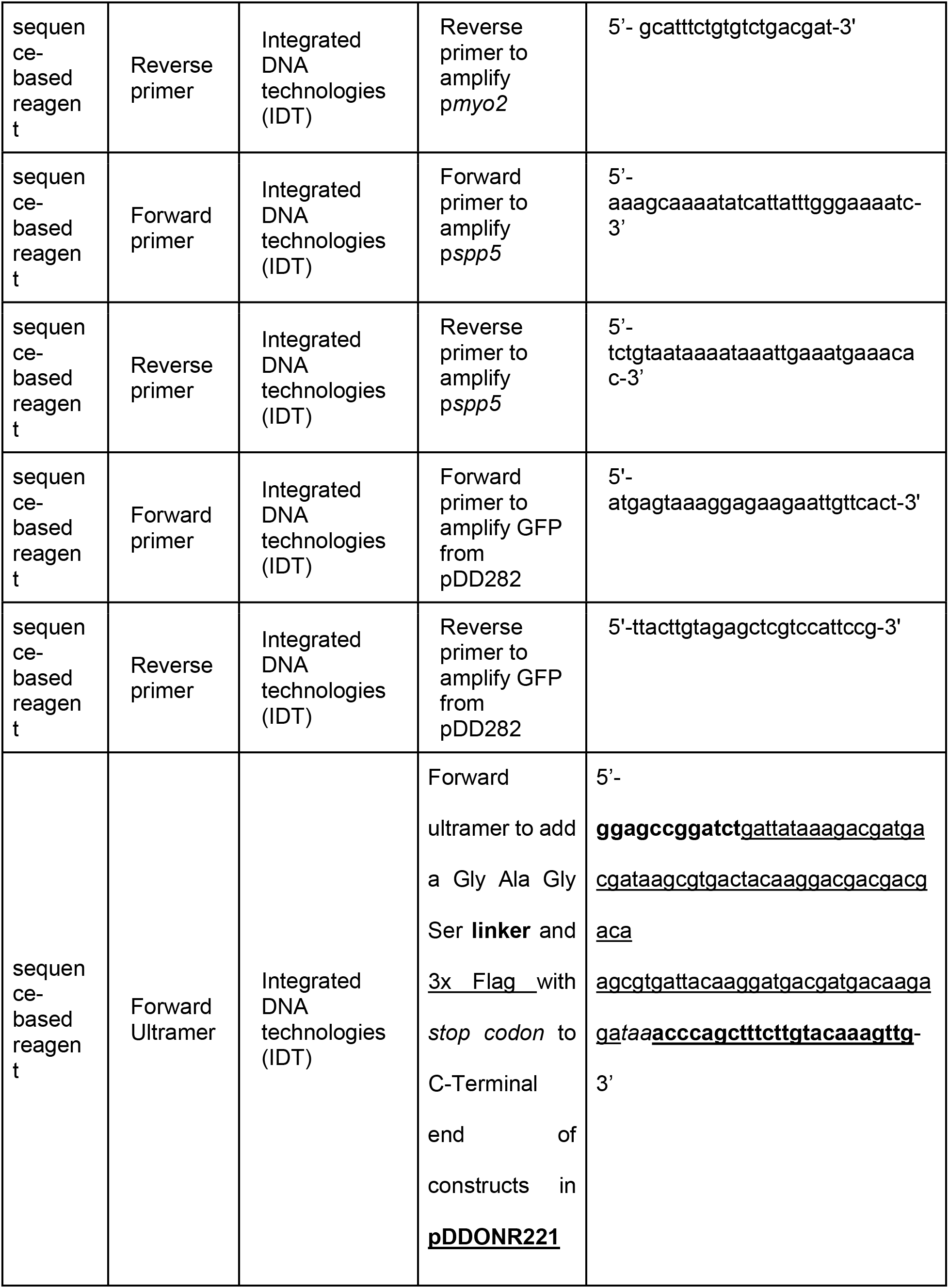
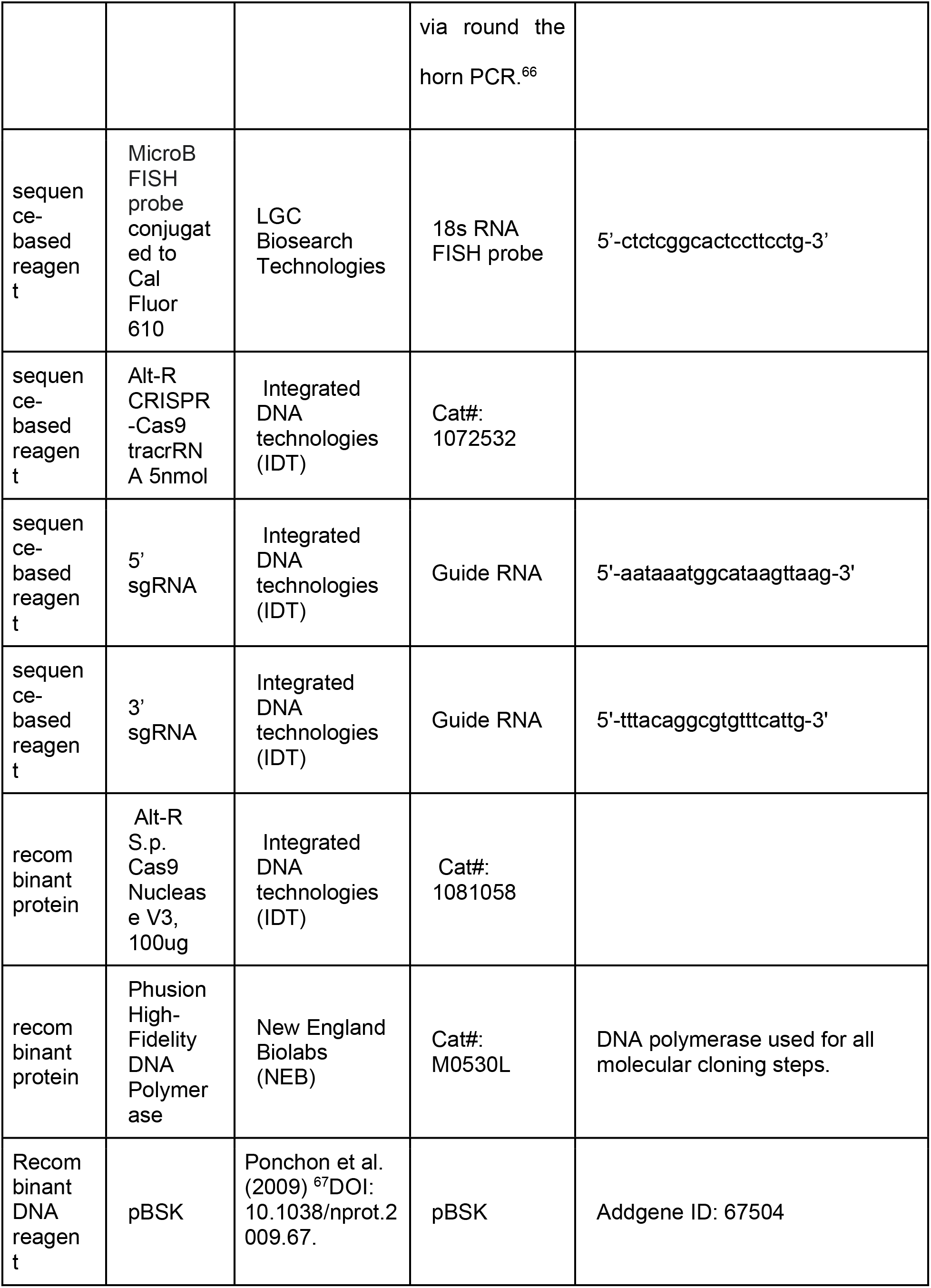
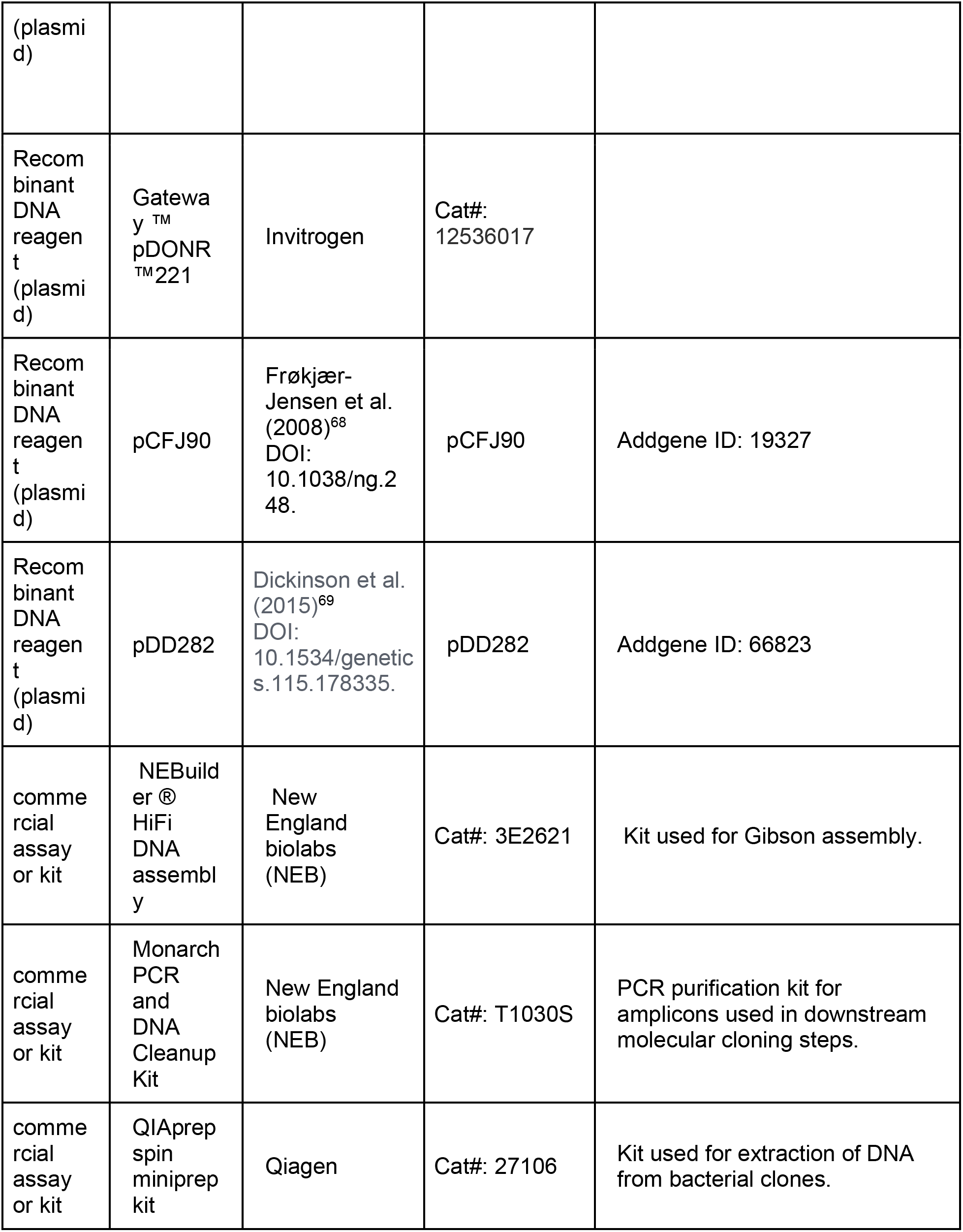
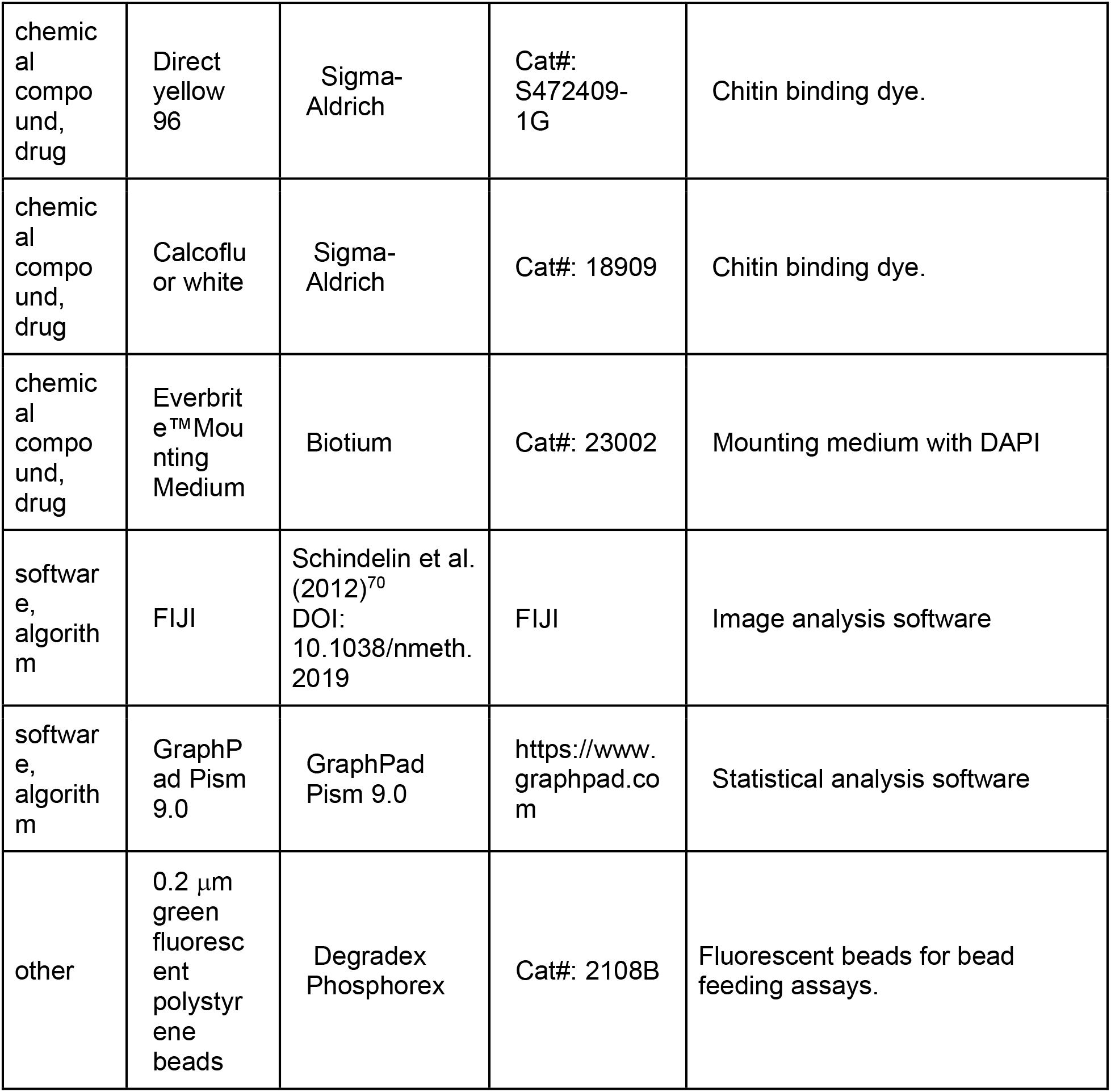

### Strain maintenance

*C. elegans* strains were grown at 21°C on nematode growth media (NGM) plates seeded with 10x saturated *Escherichia coli* OP50-1.^27^ Strains used in this study are listed in the key resources table. For all infection assays, 15-20 L4 staged animals were picked onto 10cm seeded NGM plates 4 days prior to sodium hypchlorite/1M NaOH treatment. After 4 days, heavily populated non-starved plates were washed off with 1 ml M9, treated twice with 1 ml of sodium hypochlorite/1M NaOH solution, and washed three times in 1 ml M9. Embryos were then resuspended in 5 ml of M9 and left to rock overnight at 21°C. L1s were used in subsequent experiments no later than 20 hours after bleach treatment. All centrifugation steps with live animals/embryos were performed in microcentrifuge tubes at 845xg for 30s.

Throughout the paper, L1 refers to the stage immediately post hatching or bleach synchronization, L3 refers to 24 hours and L4 refers to 48 hours post plating of bleach synchronized L1s at 21°C. L3 and L4 stage animals were washed off plates in M9 + 0.1% Tween-20, followed by an additional wash to remove residual bacteria before infection with microsporidia, or plating on PA14.

### Forward Genetic Screen

6,000 L4 N2 hermaphrodites were mutagenized with a combination of 50 mM EMS and 85.4 mM ENU for 4 hours to achieve a large diversity of mutations within the genome.^71^ P0 animals were then split and placed onto 48 10cm NGM plates, F1s bleached and resulting F2s pooled onto 5 separate plates. 180,000 L1 F2 animals were plated onto a 10 cm plate with 10 million *N. parisii* spores and 1 ml 10x saturated OP50-1. Animals were grown for 72 hours, to select for animals that display a fitness advantage phenotype with respect to N2. Each population was bleached and grown in the absence of infection for one generation, in order to prevent the effects of intergenerational immunity^27^. Two more cycles of infection followed by growing worms in the absence of infection was performed. Populations of bleached L1s were then infected with either 20 or 40 million spores and grown for 76 hours. Worms were then washed into 1.5 ml microcentrifuge tubes and 1 ml of stain solution (1x PBS/0.1% Tween-20/2.5 mg/ml DY96/1% SDS) was added. Samples were incubated with rotation for 3.5 hours and then washed 3 times with M9 + 0.1% Tween-20. Individual worms that had embryos, but not spores, were picked to individual plates. Each of the four *fawn* strains was isolated from a different mutant pool.

### Whole genome sequencing

N2 and *fawn* isolates were each grown on a 10 cm plate until all *E. coli* was consumed. Each strain was washed off with M9 and frozen at -80°C. DNA was extracted using Gentra puregene Tissue Kit (QIAGEN). Samples were sequenced on an Illumina HiSeq 4000, using 100 base paired end reads.

### MIP-Map

Molecular inversion probes were used to map the underlying causal mutations in *fawn* isolates as previously described.^35^ Briefly, *fawn* hermaphrodites were crossed to males of the mapping strain DM7448 (VC20019 Ex[*pmyo3::*YFP]) hereafter referred to as VC20019. Next, 20 F1 hermaphrodite cross progeny, identified as those carrying *pmyo3*::YFP, were isolated and allowed to self. F2s were then bleached, and 2,500 L1s were exposed to a medium-2 dose of *N. parisii* spores representing the first round of selection. Two plates of 2,500 F3 L1s were set up. The experimental plate was grown in the absence of infection for one generation, to negate intergenerational immunity.^27^ A second plate of 2,500 L1s was allowed to grow for 72 hours and then frozen in H_2_O at -80°C, until used for genomic preparation. The selection and rest steps were repeated once more, and a second frozen sample of worms was taken at the end of the mapping experiment. This process was also performed for a cross between N2 hermaphrodites and males of the mapping strain VC20019, as a negative control to identify non-causal loci that may be selected for reasons other than resistance to infection. Two genomic preparations, corresponding to the two rounds of selection, were used as template for MIP capture, to generate multiplexed libraries for sequencing. An Illumina Mini-seq was used to generate sequencing data that was subjected to demultiplexing via R, and selection intervals were defined as those immediately adjacent to the region on the chromosome carrying the fewest proportion of reads corresponding to the mapping strain, VC20019. This interval was then used to scan for putative causal alleles, resulting in the identification of the four *aaim-1* alleles in the four *fawn* isolates.

### Identification of causal gene

Variants were identified using a BWA-GATK pipeline. Briefly, sequencing reads were checked for sequence quality using FastQC (http://www.bioinformatics.babraham.ac.uk/projects/fastqc/) and bases lower than a quality threshold of 30 were trimmed off with Trimmomatic using a sliding window of 4 bases and minimum length of 36 bases.^72^ Reads were aligned to the *C. elegans* N2 reference genome (release W220) using BWA-mem.^73^ Alignments were sorted by coordinate order and duplicate reads removed using Picard (https://github.com/broadinstitute/picard). Prior to variant calling, reads were processed in Genome Analysis Tool Kit (GATK) v3.8.1,^74^ to perform indel realignment and base quality score recalibration using known *C. elegans* variants from dbSNP, build 138 (http://www.ncbi.nlm.nih.gov/SNP/). GATK HaplotypeCaller was used to call variants, and results were filtered for a phred-scaled Qscore > 30 and to remove common variants found previously in multiple independent studies. Finally, Annovar ^75^ was used to obtain a list of annotated exonic variants for each sequenced strain.

### Microsporidia infection assays

*N. parisii* (ERTm1) and *N. aususbeli* (ERTm2) spores were prepared as described previously.^27^ All infections were carried out on 6-cm NGM plates, unless otherwise specified by spore dose (see Supplemental table 1), or experimental method. 1,000 bleach-synchronized L1s were added into a microcentrifuge tube containing 400 μl of 10X *E. coli* OP50-1, and spores. After pipetting up and down, this mixture was top plated onto an unseeded 6-cm NGM plate, and left to dry in a clean cabinet, prior to incubation at 21°C for 72 hours. Infections set up on 3.5-cm plates used 160 μl of 10x *E. coli* OP50-1 and 400 L1s.

### Infection of embryos hatched on plates

Twenty-five 72-hour old synchronized animals of each strain were picked onto 3.5-cm unseeded NGM plates seeded with 16 μl of 10x *E. coli* OP50-1. Plates were incubated at 21°C for two hours. Adults were then picked off, and a mixture of 144 μl of 10x *E. coli* OP50-1 and a low dose of *N. parisii* spores were added to each plate. Animals were fixed and stained after 72 hours.

### Pulse-chase infection assay

6,000 bleach synchronized animals were exposed to a medium-1 (Figure S4) or medium-3 (Figure 3) dose of spores, 10 μl of 10x *E. coli* OP50-1 in a total volume of 400 μl made up with M9. To assay pathogen clearance 3 hpi, animals were washed off in 1 ml M9 + 0.1%Tween-20, and split into two populations. The first half was fixed with acetone to represent initial infectious load, while the other half was washed twice in M9 + 0.1% Tween-20 to remove residual spores in the supernatant and prevent additional infection from occurring. These washed worms were then plated on 6-cm unseeded NGM plates with 40 μl 10x OP50-1, and 360 μl M9 and left to incubate at 21°C for 21 additional hours before fixation.

### Spore localization and firing assays

Strains were infected as described for the pulse infection assays for either 45 minutes or 3 hours. Animals were then washed off plates, fixed, and stained with DY96 and an *N. parisii* 18S RNA FISH probe. FISH^+^ DY96^+^ events represent unfired spores, FISH^−^ DY96^+^ events represent fired spores, and FISH^+^ DY96^-^ events represent sporoplasms. Percentage of fired spores is defined as the number of FISH^−^ DY96^+^ events over the total number of spores.

To assess spore orientation, the localization of Calcofluor White spores relative to the apical membrane of the apical intestine was measured in live anaesthetized animals using differential interference contrast microscopy. To determine if a spore was angled, straight lines were extended from both ends of the spore independently. If either of these two lines crossed the apical membrane, a spore was considered angled. If not, the spore was considered parallel. Distance of spores from the apical membrane was assessed by measuring perpendicular distance from the central edge of a parallel spore to the apical membrane. All measurements were performed with FIJI^70^ using the angle tool or the straight line tool respectively, followed by the Analyze → measure option. Images in Figure 4a were taken in N2 and *aaim-1 (kea22)* animals carrying PGP-1::GFP to label the apical intestinal membrane, thus outlining the lumen.

### Intestinal lumen measurements

Measurements were performed on live anaesthetized worms used for spore localization assays (see above). The width of the lumen was determined by extending a straight line from the apical membrane on one end of the worm to that directly across on the other end, at the midpoint of the intestine, and the distance measured in FIJI, via the straight line tool followed by the Analyze → measure option.

### Fixation

Worms were washed off infection plates with 700 μl M9 +0.1% Tween-20 and washed once in 1 ml M9+0.1%Tween-20. All microsporidia infected samples were fixed in 700 μl of acetone for 2 minutes at room temperature prior to staining. All *P. aeruginosa* PA14::DsRed infected samples, as well as competitive fitness assays involving RFP::ZNFX-1 were fixed in 500 μl of 4% paraformaldehyde (PFA) for 30 minutes at room temperature prior to mounting on slides.

### Live imaging

Animals were mounted on 2% Agarose pads in 10 μl of 10-25mM Sodium Azide. This technique was used for spore localization assays, transcriptional reporter imaging, and assessing PA14::DsRed colonization in transgenic animals.

### Chitin Staining

The chitin binding dye Direct yellow 96 (DY96) was used to assess host fitness (gravidity) as well as parasite burden. 500 μl of DY96 solution (1 x PBST, 0.1% SDS, 20 μg/ml DY96) was added to washed worm pellets and left to rock for 20-30 minutes at room temperature. Worms were then resuspended in 20 μl of EverBrite™ Mounting Medium (Biotium), and 10 μl mounted on glass slides for imaging.

To prestain spores prior to infection, 0.5 μl of Calcofluor white solution (CFW) (Sigma- Aldrich 18909) was added per 50 μl of spores, pipetted up and down gently and left for 2 minutes at room temperature prior to infection.

### FISH staining

To quantify the number of sporoplasms in *N. parisii* infected animals, the MicroB FISH probe (ctctcggcactccttcctg) labelling *N. parisii* 18S RNA was used. Animals were fixed in acetone, washed twice in 1 ml PBST, and once in 1 ml of hybridization buffer (0.01% SDS, 900 mM NaCl, 20 mM TRIS pH 8.0). Samples were then incubated overnight in the dark at 46 °C with 100 μl of hybridization buffer containing 5 ng/μl of the MicroB FISH probe conjugated to Cal Fluor 610 (LGC Biosearch Technologies). Samples were then washed in 1ml of wash buffer (Hybridization buffer + 5 mM EDTA), followed by incubation with 500 μl wash buffer at 46 °C in the dark. To visualize sporoplasms and spores simultaneously, the final incubation was replaced with 500 μl DY96 solution and incubated in the dark at room temperature prior to resuspension in 20 μl of EverBrite™ Mounting Medium (Biotium).

### Microscopy and image quantification

All imaging was performed using an Axio Imager.M2 (Zeiss), except for images of the transcriptional reporter in Figure S6, which were generated using an Axio Zoom V.16 (Zeiss) at a magnification of 45.5x. Images were captured via Zen software and quantified under identical exposure times per experiment. Gravidity is defined as the presence of at least one embryo per worm, and animals were considered infected by 72 hours if clumps of spores were visible in the body of animals as seen by DY96. FISH-stained animals were considered infected if at least one sporoplasm was visible in intestinal cells.

To quantify fluorescence within animals (Pathogen burden, bead accumulation, and GFP), regions of interest were used to outline every individual worm from anterior to posterior, unless otherwise specified in the methods. Individual worm fluorescence from variable assays (GFP or DsRed) were subjected to the “threshold” followed by “measure” tools in FIJI.^70^ To assess PA14::DsRed burden in transgenic animals, regions of interest were generated from the beginning of the intestines (int1) to the posterior end of the worm to prevent the p*myo2*::mCherry co-injection marker signal from interfering with quantifications. When assessing pathogen burden in gravid animals stained with DY96, thresholding was used to quantify spore signal without including signal from embryos.

### Pseudomonas aeruginosa infection experiments

For all *Pseudomonas* assays, a single colony was picked into 3 ml of LB and grown overnight at 37°C, 220 rpm for 16-18 hours. 10 μl (for 3.5-cm plate) or 50 μl (for 6-cm plate) of culture was spread onto slow killing (SK) plates to form a full lawn, except in the case of competitive fitness assays (see below). Seeded plates were placed at 37°C for 24 hours, followed by 25°C for 24 hours prior to use. Plates were seeded fresh prior to each experiment. To assess colonization, 1,000 synchronized animals were grown on PA14::DsRed for either 24 or 48 hours at 25°C. Animals were washed off with 1 ml M9+ 0.1% Tween-20, and washed twice thereafter, prior to fixation.

To quantify survival of individual strains on PA14, 3.5-cm SK plates were seeded with 10 ul of PA14::DsRed, to form full lawns. 60 L4s were picked onto each of three, 3.5-cm plates per strain, and 24 hours later, 30 animals from each were picked onto a new 3.5-cm plate (T24hrs). Survival was monitored from 24 hours post L4, three times per day. Survival was assessed based on response to touch. Carcasses were removed, and surviving animals were placed onto fresh 3.5-cm plates every 24 hours. Animals were grown at 25°C for the duration of the experiment. Technical triplicate data was pooled to represent a single biological replicate. The experiment was carried out until no more worms had survived. Survival curves were generated via GraphPad Prism 9.0, and the Log rank (mantel-cox) test was used to generate P-values. TD_50_ values were calculated as previously described,^41^ utilizing GraphPad Prism 9.0 and applying a non-linear regression analysis on survival curves.

### Staphylococcus aureus infection experiments

3.5 cm Tryptic soy agar (TSA) plates supplemented with 10ug/ml Nalidixic acid (Nal) and seeded with *S. aureus* NCTC8325 were utilized and survival quantified as described previously.^76^ Briefly, a 1:10 dilution of an overnight *S. aureus* culture was utilized to seed 3.5 cm TSA + Nal plates, incubated at 37°C for 3 hours and stored at 4°C overnight. 30 L4s were picked onto three TSA+Nal plates per strain, and survival quantified three times a day until all animals were dead. Animals were transferred to new seeded TSA + Nal plates every 24 hours. Survival was assessed based on response to touch. Technical triplicate data was pooled to represent a single biological replicate. Survival curves were generated via GraphPad Prism 9.0, and the Log rank (mantel-cox) test was used to generate P-values.

### Transgenic strain construction

N2 or *aaim-1 (kea22)* animals were injected with a 100 ng/μl injection mix composed of 50 ng/μl of template, 5 ng/μl of p*myo2*::mCherry, and 45 ng/μl of pBSK. Three independent lines were generated for each injected construct.

Gateway BP cloning^77, 78^ was performed to insert AAIM-1 and GFP into pDONR221. Around the horn PCR,^66^ was used to insert a 3x Flag sequence at the C-terminus of this construct. Gibson assembly was used to generate different tissue specific clones driving *aaim-1* expression. P*aaim- 1, aaim-1* and p*spp-5* were cloned from N2 genomic DNA, *pmyo2* was cloned from pCFJ90. GFP and 3x Flag sequences were cloned from pDD282. *SPΔaaim-1*was amplified from *aaim-1*::3xFlag in pDONR221 by omitting the first 17 amino acids, the putative secretion signal as predicted via SignalP 5.0.^36^ All clones possessed an *unc-54* 3’ UTR. See key resources table for primer sequences.

### CRISPR-Cas9 mutagenesis

To generate a deletion allele of *aaim-1* via CRISPR-Cas9 mutagenesis, steps were taken as described here.^79^ Briefly, 2 crRNA’s were designed using CRISPOR,^80^ near the start and stop sites of *aaim-1* and generated via IDT. A repair template was designed to contain 35 base pairs of homology upstream and downstream of the cut sites. *Streptococcus pyogenes* Cas9 3NLS (10ug/ul) IDT and tracrRNA (IDT #1072532) were utilized. Reaction mixes were prepared as described previously.^79^ pRF4^81^ was co-injected with the Cas9 ribonucleoprotein, and F1 rollers picked. Deletions were identified via PCR primers situated outside the cut sites.

### Bead-feeding assays

1,000 synchronized L1 animals were mixed with 0.2 μm green fluorescent polystyrene beads (Degradex Phosphorex) at a ratio of 25:1 in a final volume of 400 μl containing 10 μl of 10x *E. coli* OP50-1, 16 μl of beads and up to 400 μl with M9. Animals were incubated with beads for 3 hours, washed off with M9 + 0.1% Tween-20 and fixed with 4% PFA for 30 min at room temperature. Bead accumulation was measured as a percentage of the total animal exhibiting fluorescent signal, using FIJI.

### Lifespan Assays

Lifespan assays were performed as described previously.^82^ In brief, 120 synchronized L4 animals were utilized per strain, with every 15 animals placed on a single 3.5-cm NGM plate (A total of 8 plates, with 15 animals each per strain). Animals were transferred to a new seeded 3.5-cm NGM plate every 2 days, for a total of 8 days (4 transfers), ensuring no progeny were transferred alongside adults. After day 8, survival was quantified daily, on the same plate, via response to touch. Any animals that exhibited internal hatching, protruding intestines, or were found desiccated on the edges of the plate were censored. Survival curves were generated via GraphPad Prism 9, and the Log rank (mantel-cox) test was used to generate P values.

### Immunofluorescence (IF)

IF was performed as described previously,^83^ however all steps post-dissection were performed in microcentrifuge tubes, and intestines were pelleted on a mini tabletop microcentrifuge for a few seconds. Briefly, animals were dissected to extrude intestinal tissue. Two 25-mm gauge needles on syringes were used to create an incision near the head and/or tail of the animals. Dissections were performed in 5 μl of 10 mM levamisole on glass slides to encourage intestinal protrusion. Fixation, permeabilization and blocking was performed as described previously.^83^ Primary M2 anti-Flag antibody (Sigma F1804) was used at 1:250 overnight at 4°C, and secondary goat anti-mouse Alexa fluor 594 (Thermo Fisher A32742) at 1:300 for 1 hour at room temperature. Animals were mounted in 20 μl of EverBrite™ Mounting Medium (Biotium) and placed on glass slides for imaging.

### Competitive fitness assays

N2 or *aaim-1* mutants were grown together with RFP::Znfx1 YY1446 *(gg634)*, which labels the germ granules and can be observed in all developmental stages^84^. For *N. parisii* infections, 10-cm NGM plates were seeded with 1 ml of 10x OP50-1 and a medium-2 dose of spores (no spores were used for uninfected plates). 10 L1s from each strain were picked onto lawns of spores and *E. coli* OP50-1 immediately after drying, and grown for 8 days at 21°C, washed off with M9 + 0.1% Tween-20, and fixed. For *P. aeruginosa* infections, 3.5-cm SK plates were seeded with a single spot of 20 μl of PA14 in the center of the plate. 10 L1s of each strain were placed on plates and grown at 21°C for 8 days and then washed off with M9 + 0.1% Tween-20. The percentage of animals that did not display RFP germ granules (i.e. N2 or *aaim-1* mutants) was determined by quantifying all animals on the plate, including F1 adults and L1/L2 stage F2 animals.

### Co- infections with N. parisii and P. aeruginosa

Co-infection assays were performed by first pulse infecting co-infection and *N. parisii* single infection groups with a maximal dose of spores for three hours on unseeded 6-cm NGM plates as described above. PA14::DsRed single infections were pulsed with a volume of M9 to match that of the spores. Animals were then washed off in 1ml of M9 + 0.1%Tween-20, followed by 2 more washes, prior to placement on full lawns of PA14::DsRed on a 6-cm SK plates prepared as described above. *N. parisii* single infections were placed on a 6-cm NGM plate pre-seeded with 200 μl of 10x OP50-1. Plates were incubated at 21°C.

### Phylogenetic analysis

Homology between AAIM-1 and other proteins was determined with protein BLAST (https://blast.ncbi.nlm.nih.gov/Blast.cgi) using default parameters. Sequences with less than E-5 were aligned using MUSCLE (https://www.ebi.ac.uk/Tools/msa/muscle/) using default parameters. Phylogenetic tree of homologs was generated using RAxML BlackBox https://raxml-ng.vital-it.ch/<u>#/</u> using default parameters and 100 boot straps. Tree was visualized using FigTree v1.4.4 (http://tree.bio.ed.ac.uk/software/figtree/).

### Statistical analysis

All data analysis was performed using GraphPad Prism 9.0. One-way Anova with post hoc (Tukey test) was used for all experiments unless otherwise specified in figure legends. Statistical significance was defined as p < 0.05.

## Acknowledgements

We thank Ashley M. Campbell, Alexandra R. Willis, and Kristina Sztanko for providing helpful comments on the manuscript. This work was supported by the Canadian Institutes of Health Research grant no. 400784 and an Alfred P. Sloan Research Fellowship FG2019-12040 (to A.W.R.). This work was supported by National Institutes of Health (www.nih.gov) under R01 AG052622 and GM114139 to E.R.T. Some strains were provided by the CGC, which is funded by NIH Office of Research Infrastructure Programs (P40 OD010440) and we thank WormBase.

## Author contributions

**H.T.E.J. and A.W.R.** designed experiments, analyzed results, and co-wrote the paper.

**H.T.E.J.** conducted all experiments, except the initial forward genetic screen performed by A.W.R.

**C.M.** designed and performed bioinformatic analysis for the MIP-map experiment.

**M.R.S.** analyzed whole genome sequencing to identify causal mutations in *fawn* animals.

**E.R.T., A.G.F., and A.W.R.** provided mentorship and acquisition of funding.

## Supplemental material

**Figure S1.**
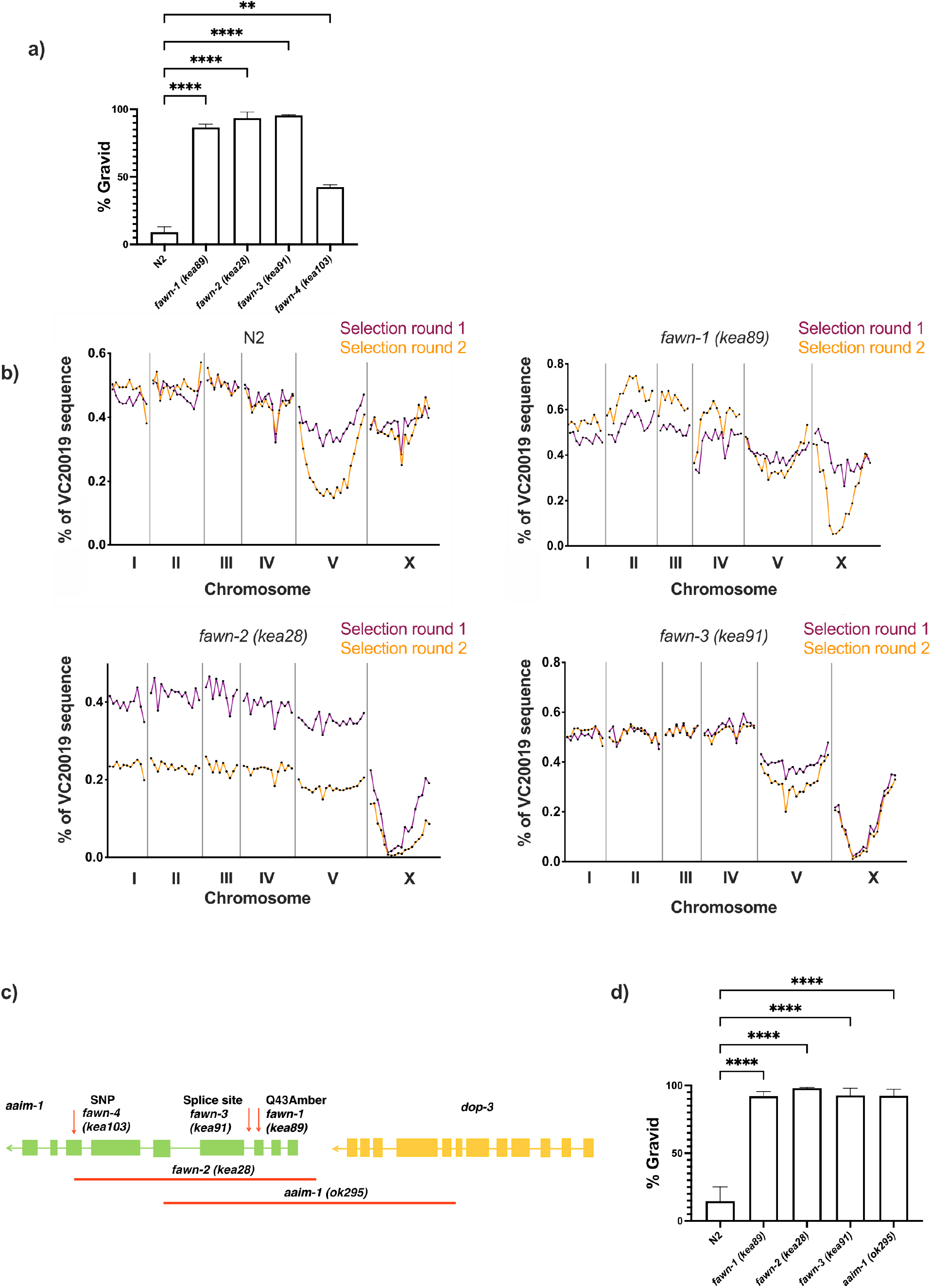
Mapping and validation of *aaim-1* as the gene associated with resistance to *N. parisii*. (a) N2 or *fawn* animals were infected with a medium-3 dose of *N. parisii* spores on 6-cm plates, fixed at 72 hours, and stained with direct-yellow 96 (DY96). Graph displays percentage of gravid worms. Data is from three independent replicates of 66-300 worms each. (b) F2 recombinants between the mapping strain VC20019 and either N2, *fawn-1*, *fawn-2*, or *fawn-3* were infected with a medium-2 dose of *N. parisii*. Two rounds of selection were performed (see methods). The percentage of sequencing reads mapping to the reference strain VC20019 are depicted on the Y-axis, and the linkage groups are depicted on the X-axis. Sequencing of MIPs resulted in capturing the identity of the genome at 89 distinct regions which are represented as points by their location along the X-axis coordinates. A significantly diminished percentage of VC20019 indicates an enrichment of non-mapping genomic sequence in that region. (c) Schematic representing the location and nature of the different *aaim-1* alleles. Boxes represent exons, and connecting lines represent introns. Arrows represent point mutations and solid red lines represent large deletions. *fawn-2 (kea28)* has a 2.2 kb deletion and *aaim-1 (kea22)* has a 2.3 kb deletion. *aaim-1 (ok295)* possesses a large deletion overlapping two different genes, *aaim-1* and *dop-3*, the boundaries of which are unclear.^41, 85^ (d) L1 stage N2 and *aaim-1* mutant animals were infected with a high dose of *N. parisii*, fixed at 72 hours, and stained with direct-yellow 96 (DY96). Graph displays percentage of gravid worms. Data is from three independent replicates of at least 100 worms each. Mean ± SEM represented by horizontal bars. P-values determined via One-way Anova with post hoc. Significance defined as ** p < 0.01, **** p < 0.0001.

**Figure S2:**
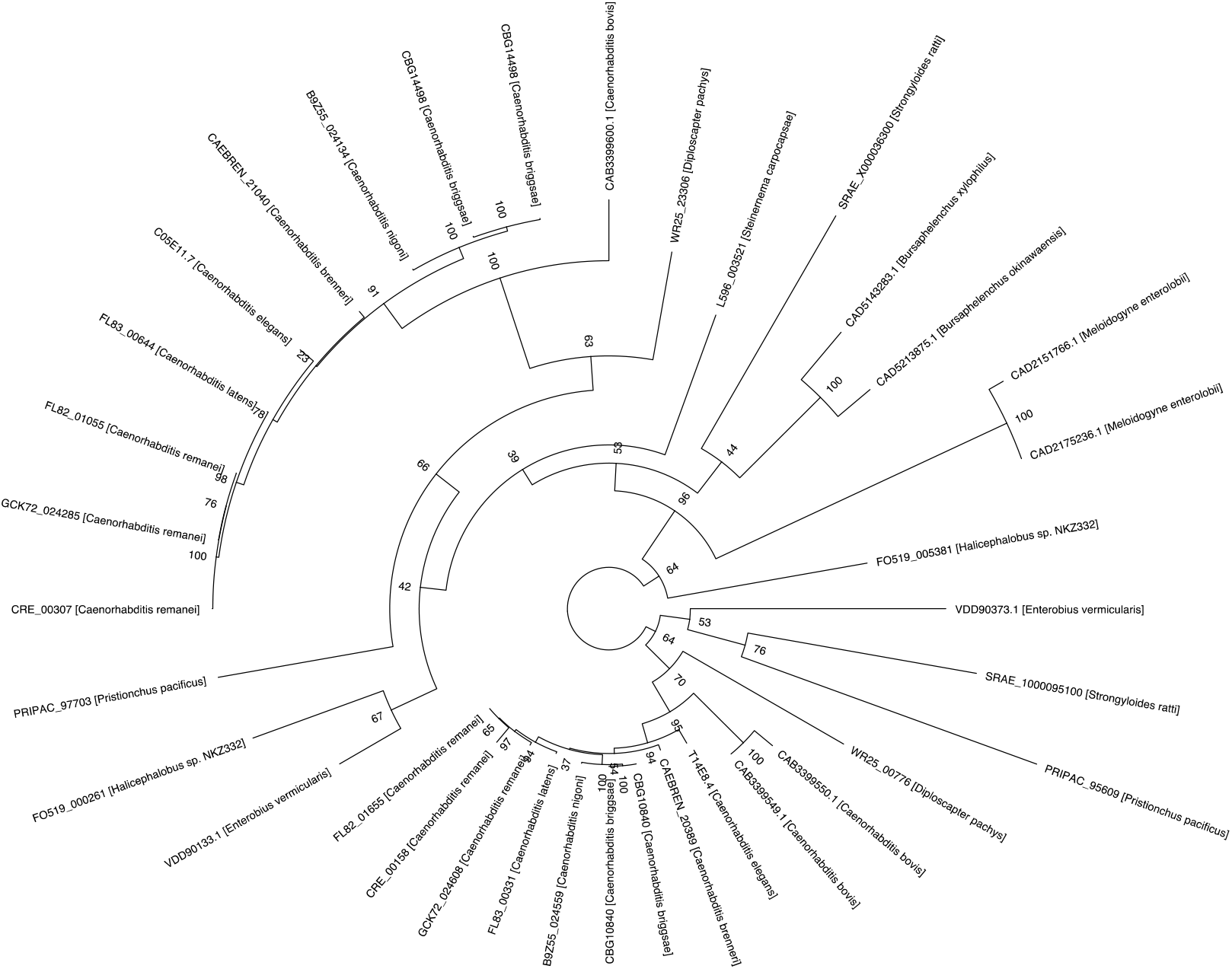
AAIM-1 is conserved in both free-living and parasitic nematodes. Phylogenetic tree of AAIM-1 homologs. Bootstrap values are shown at the nodes.

**Figure S3:**
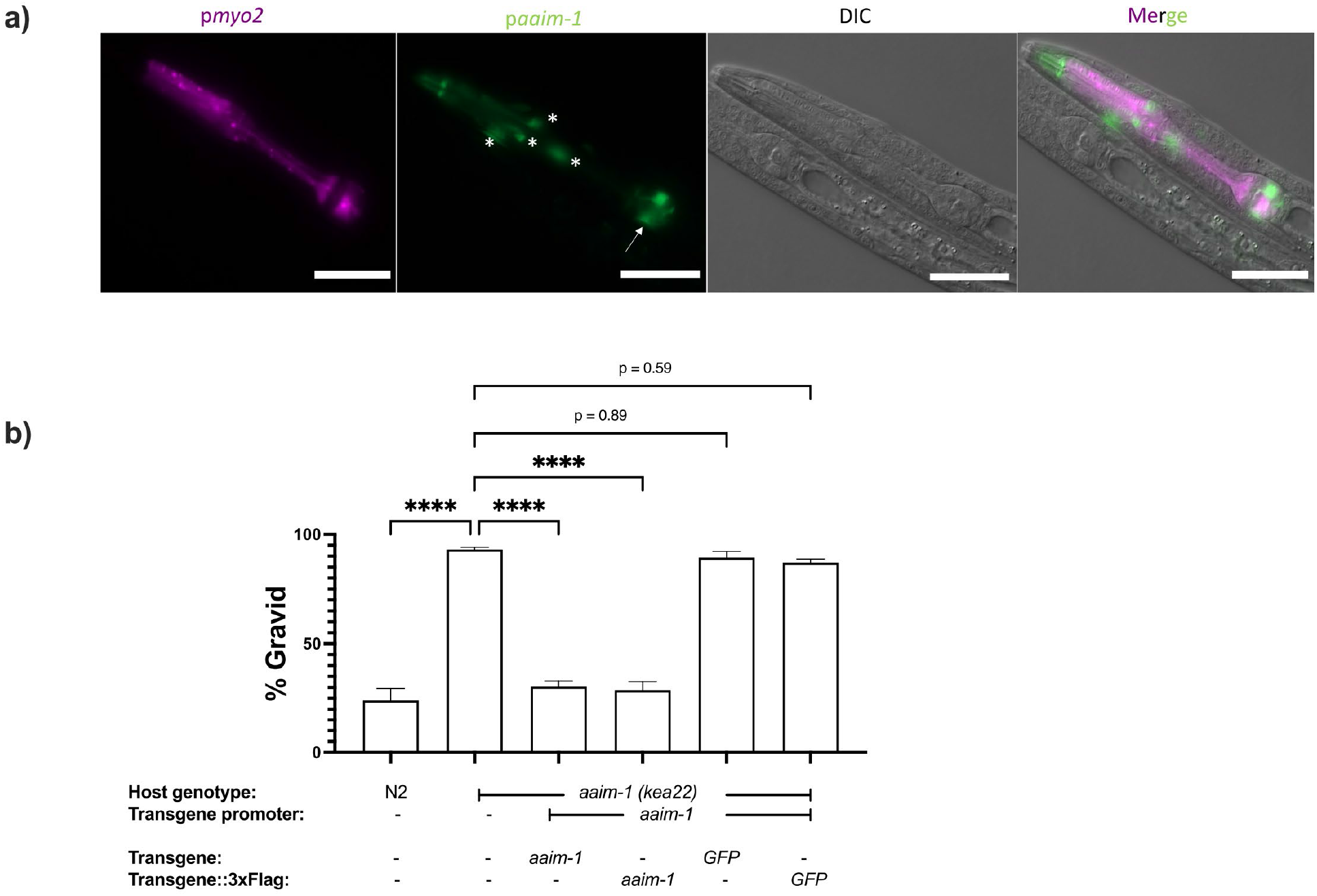
a*a*im*-1* is expressed in arcade cells and presence of C-terminal 3x Flag tag does not disrupt AAIM-1 function. (a) N2 containing an extrachromosomal array expressing GFP from the *aaim-1* promoter and mCherry (labelled in magenta) in the pharyngeal muscles were imaged at the L1 stage at 40x. Scale bar 20 μm. Arrow indicates terminal bulb, and asterisks represent arcade cells. (b) N2, *aaim- 1,* and *aaim-1* expressing extrachromosomal arrays of wild-type or 3x Flag tagged constructs were infected with a medium-2 dose of *N. parisii*, fixed at 72 hours, and stained with direct-yellow 96 (DY96). Graph displays percentage of gravid worms. Data is from three independent replicates of at least 100 worms each. Mean ± SEM represented by horizontal bars. P-values determined via one-way ANOVA with post hoc. Significance defined as **** p < 0.0001.

**Figure S4:**
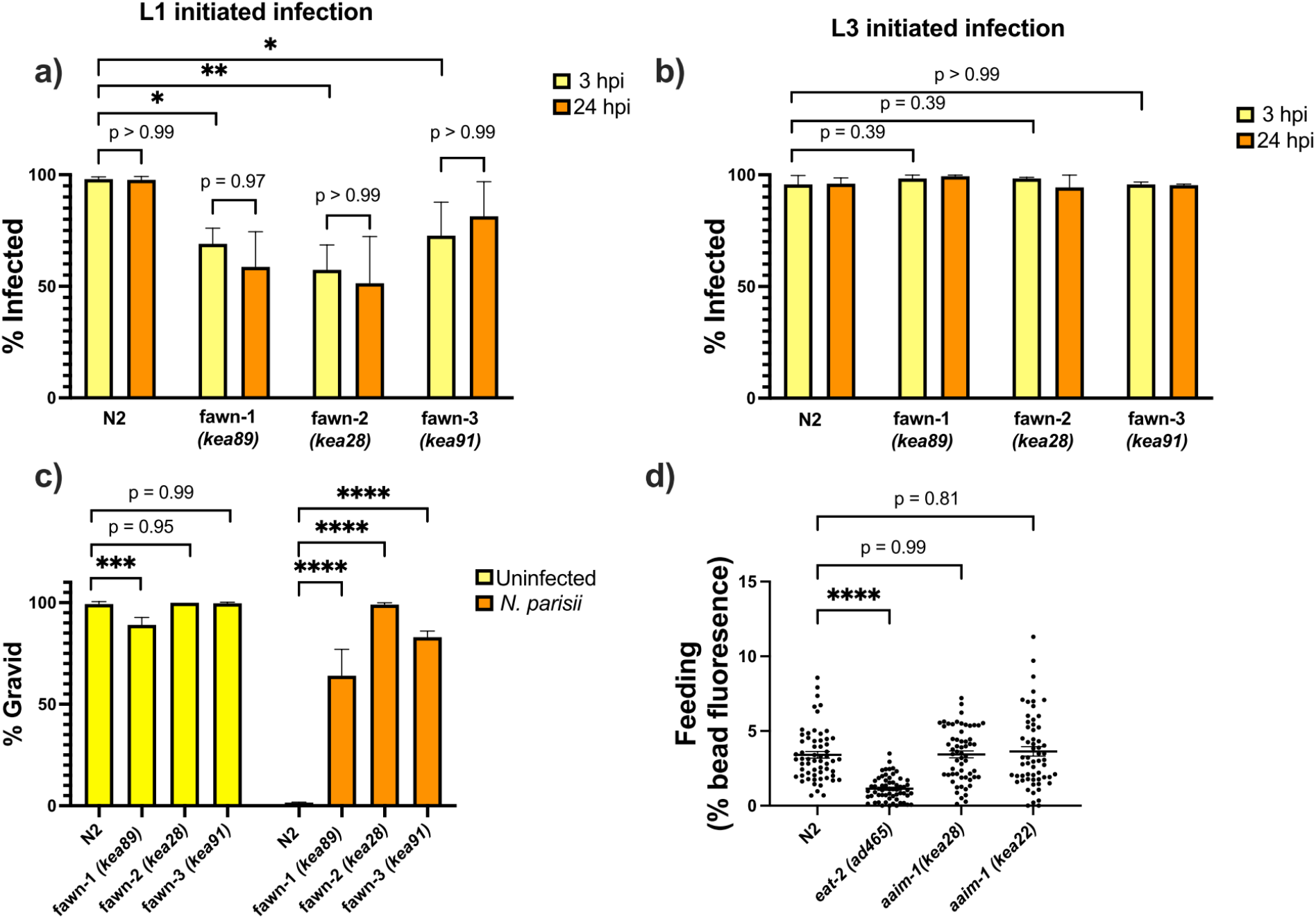
a*a*im*-1* mutants do not clear *N. parisii* and developmentally restricted *N. parisii* invasion defect is not due to a feeding defect. (a-b) N2 and *aaim-1* mutants were infected at either the L1 stage (a) or the L3 stage (b) with a medium-1 dose of *N. parisii* spores for 3 hours. Animals were then washed to remove spores and re-plated for an additional 21 hours. Worms were fixed at both the 3 hour and 24 hour timepoints and stained with an *N. parisii* 18S RNA fish probe. Worms containing either sporoplasm or meronts were counted as infected. (c) N2 and *aaim-1* adults were allowed to lay embryos on plates. Adults were removed and a low dose of *N. parisii* was added to the plate. Animals were fixed at 72 hours and stained with direct-yellow 96 (DY96). Graph displays percentage of gravid worms. (d) N2 and *aaim-1* mutants were fed fluorescent beads for 3 hours. Quantification of percentage of worm with bead fluorescence. Data is from three independent replicates of at least 100 worms each (a-c) or 20-30 worms each (d). Mean ± SEM represented by horizontal bars. P-values determined via one-way ANOVA with post hoc. Significance defined as * p < 0.05, ** p < 0.01, *** p < 0.001, **** p < 0.0001.

**Figure S5:**
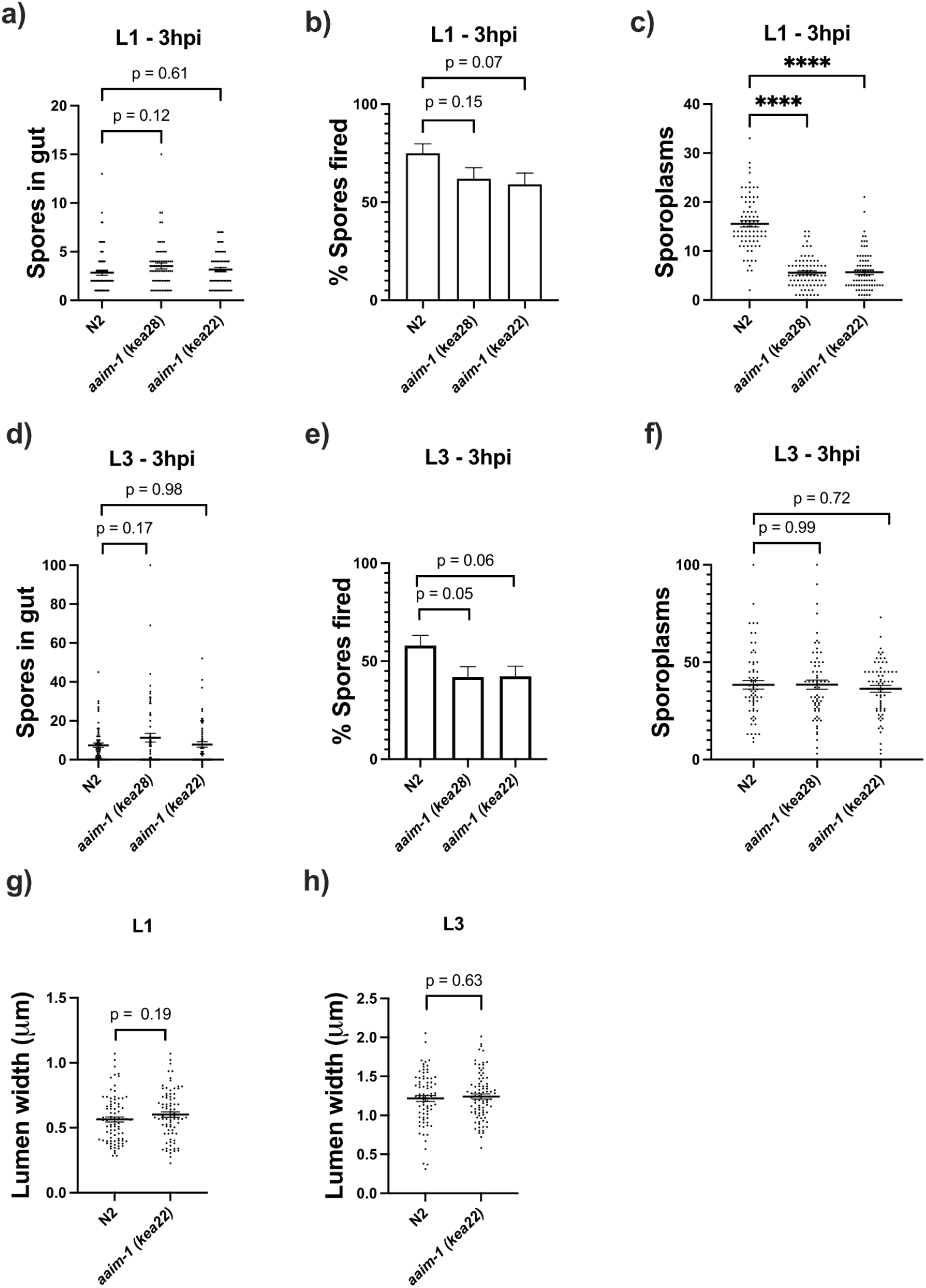
Invasion defects in *aaim-1* only occur at the L1 stage of development and a mutation in *aaim-1* does not alter the width of the intestinal lumen. (a-f) N2 and *aaim-1* animals were infected for 3 hours at L1 (a-c) or L3 (d-f), fixed, and then stained with DY96 and an *N. parisii* 18S RNA fish probe. The number of spores per animal (a,d) the percentage of spores fired (b,e) and the number of sporoplasm per worm (c,f) are displayed. (g,h) The width of the intestinal lumen was measured in L1 (g) or L3 (h) wild-type or *aaim-1* animals. (a-h) Data is from three independent replicates of 16-30 worms each. Mean ± SEM represented by horizontal bars. P-values determined via one-way ANOVA with post hoc (a-f) or Unpaired Student’s t-test (g,h). Significance defined as **** p < 0.0001.

**Figure S6:**
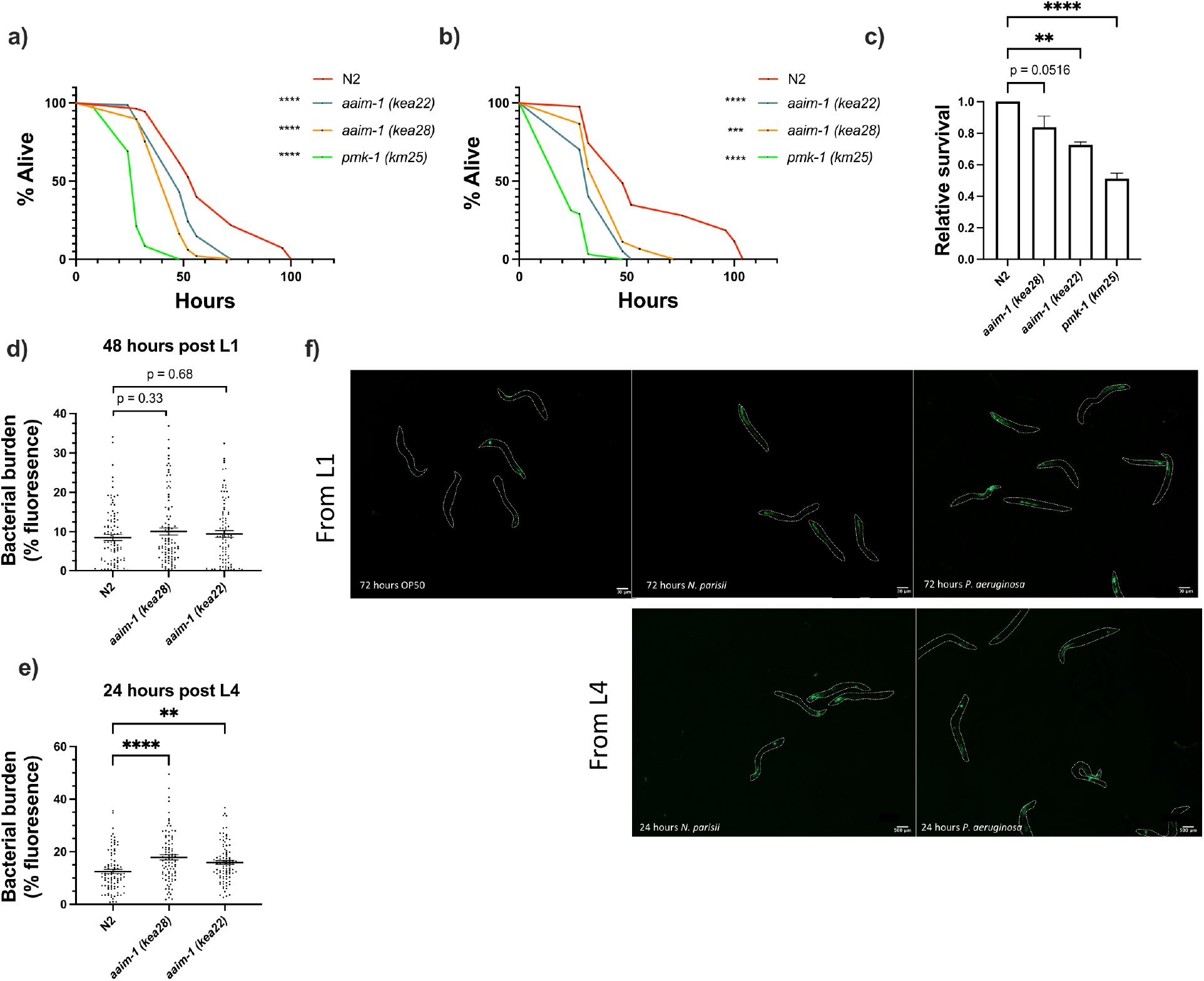
Susceptibility to *P. aeruginosa* PA14 appears at L4. (a,b) Additional replicates of survival assays of animals grown on full lawns of PA14::DsRed as in Figure 5a. (a) TD_50_ : N2 51 hours, aaim-1 *(kea28)* 46 hours, *aaim-1 (kea22)* 38 hours, and *pmk- 1 (km25)* 25 hours. (b) TD_50_ : N2 43 hours, aaim-1 *(kea28)* 30 hours, *aaim-1 (kea22)* 32 hours, and *pmk-1 (km25)* 20 hours. (c) Relative survival of mutants to N2 as calculated by mean strain TD_50_/mean N2 TD_50_. (d-e) N2 and *aaim-1* mutants were grown on PA14::DsRed 48 hours post L1 (d) or 24 hours post L4 (e). Data is from three independent replicates of 20-30 worms each. Every point represents a single worm. Bacterial burden was measured as the percentage of the animal containing PA14::DsRed via FIJI. Mean ± SEM represented by horizontal bars. (f) p*aaim- 1*::GFP::3xFlag were exposed to either PA14 or *N. parisii* 72 hours post L1 or 24 hours post L4. Animals were imaged at 45.5x, scale bar 500 μm. P-values determined via one-way ANOVA with post hoc. Significance defined as ** p < 0.01, **** p < 0.0001.

**Figure S7:**
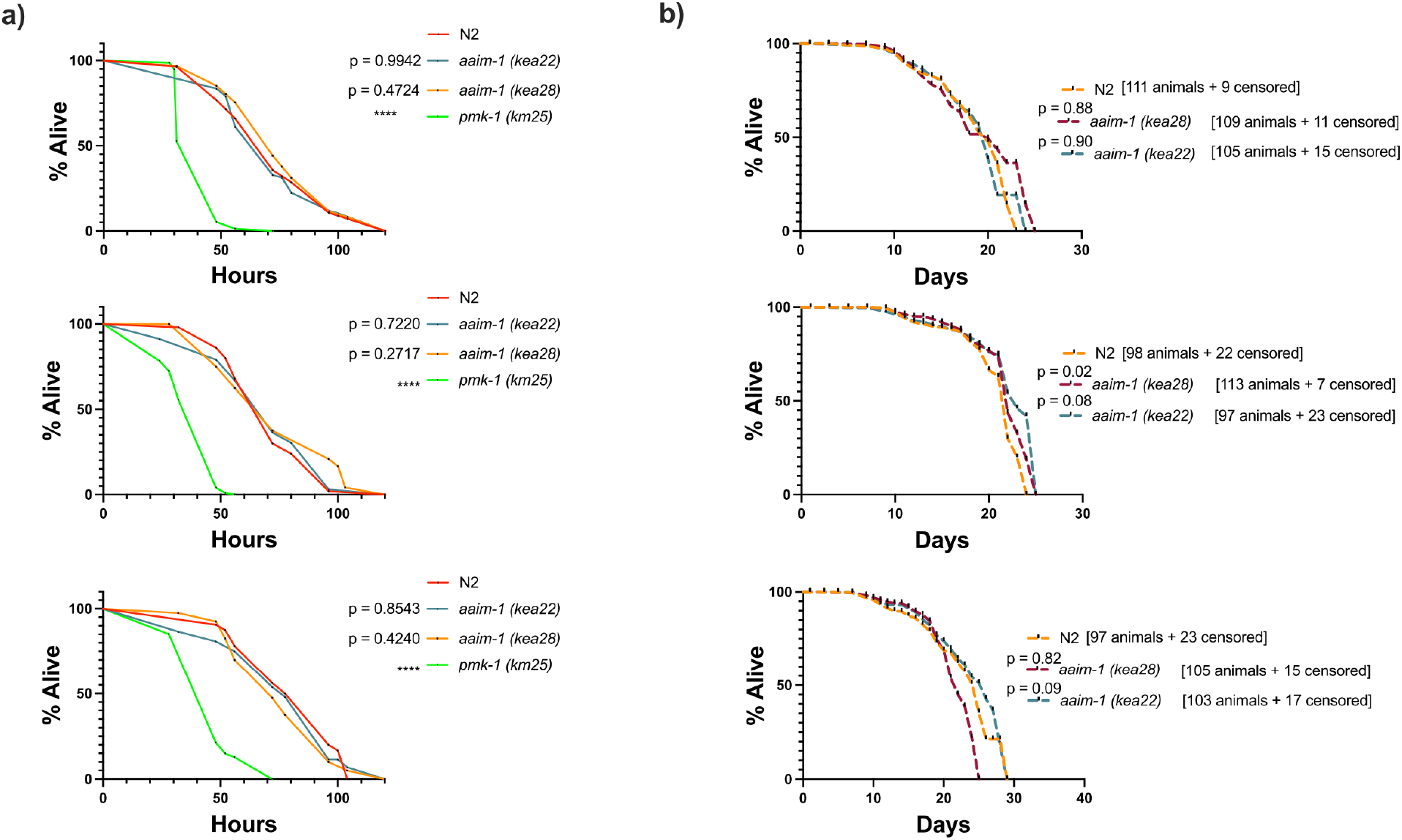
A mutation in *aaim-1* does not influence *C. elegans* defense against *S. aureus* or lifespan. (a) L4 stage N2 and *aaim-1* were plated on full lawns of *S. aureus* NCTC8325 and the percentage of animals alive was counted over the course of 120 hours. Three independent replicates are displayed. At least 40 worms were quantified per strain. (b) N2 and *aaim-1* mutants were grown on *E. coli* OP50-1 for one month, and survival measured as number of animals responsive to touch. The number of animals quantified, as well as those censored are denoted on the graph. Three independent survival assays are displayed. P-values determined via Log-rank (Mantel-Cox) test. Significance defined as **** p < 0.001.

**Figure S8:**
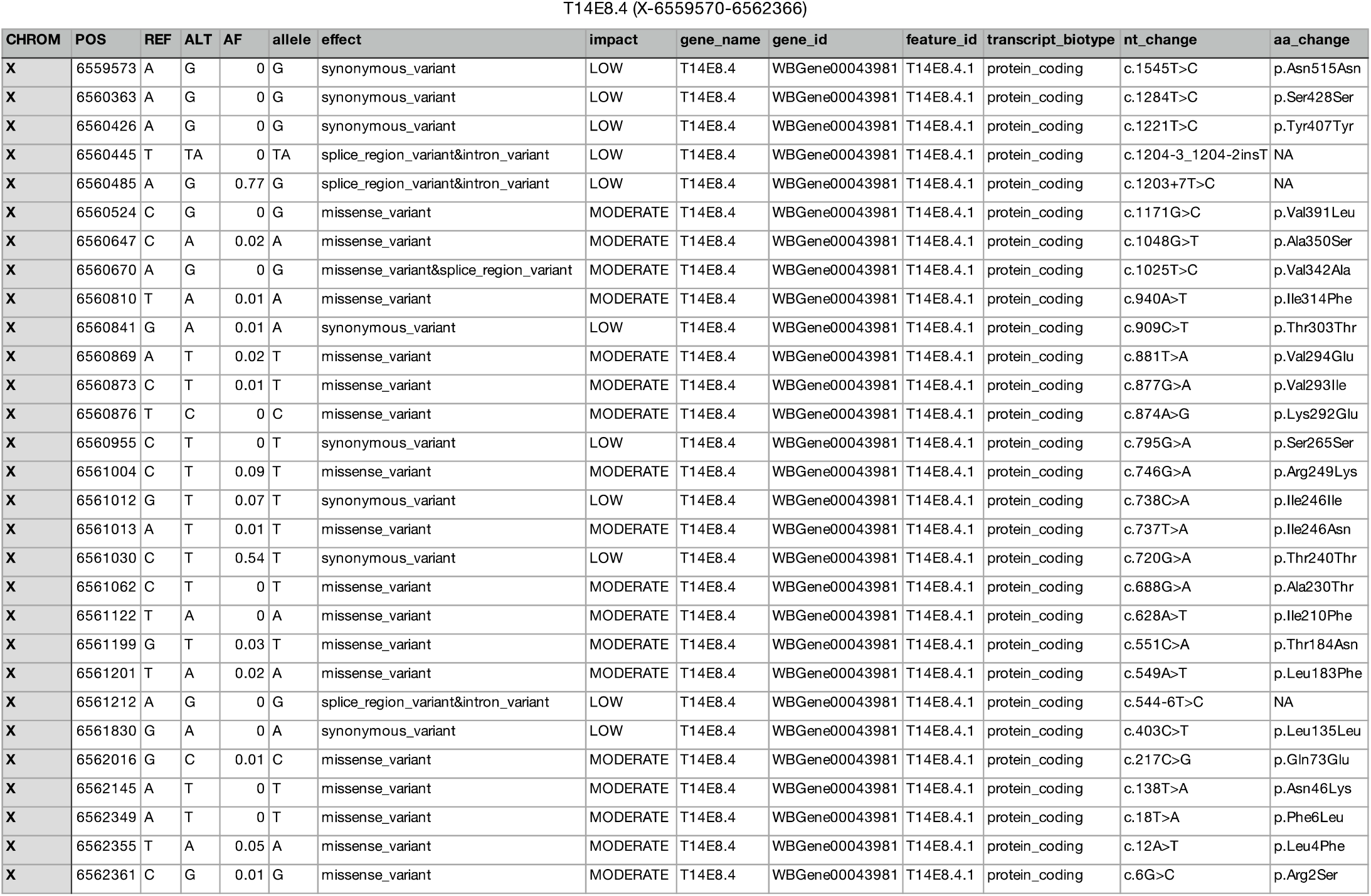
List of naturally occurring *aaim-1(T14E8.4)* variants in wild isolates of *C. elegans*. This table represents a list of *aaim-1* coding variants found to naturally occur in wild isolates of *C. elegans* generated by the CeNDR variant browser.^45^ The reference allele (REF) as well as the alternate variant (ALT) and the allele frequency (AF) are displayed for various sites (POS) across *aaim-1*. The nature (effect) and impact of these variants are depicted as well as the nucleotide changes (nt_change) and the corresponding amino acid change (aa_change). *aaim-1* does not possess any variants predicted to have a high impact, implying that there are no obvious loss of function alleles and that its retention in the wild is advantageous.

**Supplemental table 1:**
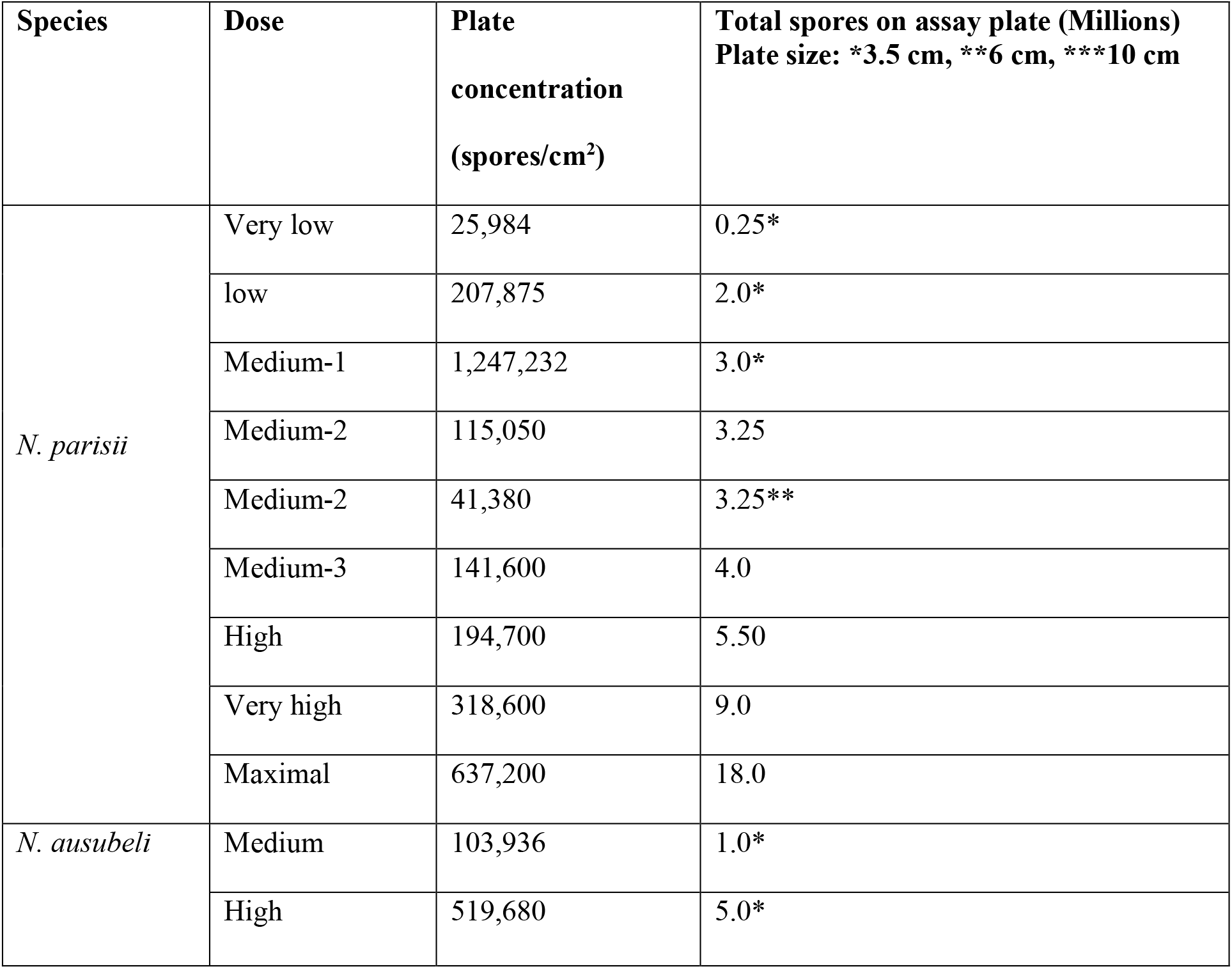
Spore doses utilized in this study.

| Species | Dose | Plate concentration<br>(spores/cm<sup>2</sup>) | Total spores on assay plate (Millions)<br>Plate size: *3.5 cm, **6 cm, ***10 cm |
| --- | --- | --- | --- |
| *N. parisii* | Very low | 25,984 | 0.25* |
|  | low | 207,875 | 2.0* |
|  | Medium-1 | 1,247,232 | 3.0* |
|  | Medium-2 | 115,050 | 3.25 |
|  | Medium-2 | 41,380 | 3.25** |
|  | Medium-3 | 141,600 | 4.0 |
|  | High | 194,700 | 5.50 |
|  | Very high | 318,600 | 9.0 |
|  | Maximal | 637,200 | 18.0 |
| *N. ausubeli* | Medium | 103,936 | 1.0* |
|  | High | 519,680 | 5.0* |

## References

1. Murareanu Brandon M. et al. Generation of a Microsporidia Species Attribute Database and Analysis of the Extensive Ecological and Phenotypic Diversity of Microsporidia. mBio 0, e01490–21.

2. Corradi, N. Microsporidia: Eukaryotic Intracellular Parasites Shaped by Gene Loss and Horizontal Gene Transfers. Annu. Rev. Microbiol. 69, 167–183 (2015).

3. Wadi, L. & Reinke, A. W. Evolution of microsporidia: An extremely successful group of eukaryotic intracellular parasites. PLoS Pathog. 16, e1008276 (2020).

4. Balla, K. M., Andersen, E. C., Kruglyak, L. & Troemel, E. R. A Wild C. Elegans Strain Has Enhanced Epithelial Immunity to a Natural Microsporidian Parasite. PLOS Pathog. 11, e1004583 (2015).

5. Routtu, J. & Ebert, D. Genetic architecture of resistance in Daphnia hosts against two species of host-specific parasites. Heredity 114, 241–248 (2015).

6. Martín-Hernández, R., et al. Nosema ceranae in Apis mellifera: a 12 years postdetection perspective. Environ. Microbiol. 20, 1302–1329 (2018).

7. Jaroenlak, P. et al. Identification, characterization and heparin binding capacity of a spore- wall, virulence protein from the shrimp microsporidian, Enterocytozoon hepatopenaei (EHP). Parasit. Vectors 11, 1–15 (2018).

8. Stentiford, G. D. et al. Microsporidia – Emergent Pathogens in the Global Food Chain. Trends Parasitol. 32, 336–348 (2016).

9. Han, B. & Weiss, L. Therapeutic targets for the treatment of microsporidiosis in humans. Expert Opin. Ther. Targets 22, (2018).

10. Han, B., Takvorian, P. M. & Weiss, L. M. Invasion of Host Cells by Microsporidia. Front. Microbiol. 11, (2020).

11. Jarkass, H. T. E. & Reinke, A. W. The ins and outs of host-microsporidia interactions during invasion, proliferation and exit. Cell. Microbiol. n/a, e13247.

12. Hayman, J. R., Hayes, S. F., Amon, J. & Nash, T. E. Developmental Expression of Two Spore Wall Proteins during Maturation of the Microsporidian Encephalitozoon intestinalis. Infect. Immun. 69, 7057–7066 (2001).

13. Hayman, J. R., Southern, T. R. & Nash, T. E. Role of Sulfated Glycans in Adherence of the Microsporidian Encephalitozoon intestinalis to Host Cells In Vitro. Infect. Immun. 73, 841– 848 (2005).

14. Southern, T. R., Jolly, C. E., Lester, M. E. & Hayman, J. R. EnP1, a Microsporidian Spore Wall Protein That Enables Spores To Adhere to and Infect Host Cells In Vitro. Eukaryot. Cell 6, 1354–1362 (2007).

15. Li, Y. et al. Identification of a novel spore wall protein (SWP26) from microsporidia Nosema bombycis. Int. J. Parasitol. 39, 391–398 (2009).

16. Wu, Z. et al. Proteomic analysis of spore wall proteins and identification of two spore wall proteins from Nosema bombycis (Microsporidia). Proteomics 8, 2447–2461 (2008).

17. Chen, L., Li, R., You, Y., Zhang, K. & Zhang, L. A Novel Spore Wall Protein from Antonospora locustae (Microsporidia: Nosematidae) Contributes to Sporulation. J. Eukaryot. Microbiol. 64, 779–791 (2017).

18. Xu, Y., Takvorian, P., Cali, A. & Weiss, L. M. Lectin Binding of the Major Polar Tube Protein (PTPl) and its Role in Invasion. J. Eukaryot. Microbiol. 50, 600–601 (2003).

19. Xu, Y., Takvorian, P. M., Cali, A., Orr, G. & Weiss, L. M. Glycosylation of the Major Polar Tube Protein of Encephalitozoon hellem, a Microsporidian Parasite That Infects Humans. Infect. Immun. 72, 6341–6350 (2004).

20. Han, B. et al. The role of microsporidian polar tube protein 4 (PTP4) in host cell infection. PLOS Pathog. 13, e1006341 (2017).

21. Han, B. et al. Microsporidia Interact with Host Cell Mitochondria via Voltage-Dependent Anion Channels Using Sporoplasm Surface Protein 1. mBio 10, (2019).

22. Luallen, R. J. et al. Discovery of a Natural Microsporidian Pathogen with a Broad Tissue Tropism in Caenorhabditis elegans. PLoS Pathog. 12, e1005724 (2016).

23. Zhang, G. et al. A Large Collection of Novel Nematode-Infecting Microsporidia and Their Diverse Interactions with Caenorhabditis elegans and Other Related Nematodes. PLOS Pathog. 12, e1006093 (2016).

24. Troemel, E. R., Félix, M.-A., Whiteman, N. K., Barrière, A. & Ausubel, F. M. Microsporidia Are Natural Intracellular Parasites of the Nematode Caenorhabditis elegans. PLoS Biol. 6, e309 (2008).

25. Troemel, E. R. New Models of Microsporidiosis: Infections in Zebrafish, C. elegans, and Honey Bee. PLOS Pathog. 7, e1001243 (2011).

26. Balla, K. M., Luallen, R. J., Bakowski, M. A. & Troemel, E. R. Cell-to-cell spread of microsporidia causes Caenorhabditis elegans organs to form syncytia. Nat. Microbiol. 1, 16144 (2016).

27. Willis, A. R. et al. A parental transcriptional response to microsporidia infection induces inherited immunity in offspring. Sci. Adv. 7, (2021).

28. Botts, M. R., Cohen, L. B., Probert, C. S., Wu, F. & Troemel, E. R. Microsporidia Intracellular Development Relies on Myc Interaction Network Transcription Factors in the Host. G3 GenesGenomesGenetics 6, 2707–2716 (2016).

29. Szumowski, S. C., Botts, M. R., Popovich, J. J., Smelkinson, M. G. & Troemel, E. R. The small GTPase RAB-11 directs polarized exocytosis of the intracellular pathogen N. parisii for fecal-oral transmission from C. elegans. Proc. Natl. Acad. Sci. 111, 8215–8220 (2014).

30. Balla, K. M., Lažetić, V. & Troemel, E. R. Natural variation in the roles of C. elegans autophagy components during microsporidia infection. PLOS ONE 14, e0216011 (2019).

31. Reddy, K. C. et al. Antagonistic paralogs control a switch between growth and pathogen resistance in C. elegans. PLoS Pathog. 15, e1007528 (2019).

32. Tecle, E. et al. The purine nucleoside phosphorylase pnp-1 regulates epithelial cell resistance to infection in C. elegans. PLOS Pathog. 17, e1009350 (2021).

33. Luallen, R. J., Bakowski, M. A. & Troemel, E. R. Characterization of Microsporidia-Induced Developmental Arrest and a Transmembrane Leucine-Rich Repeat Protein in Caenorhabditis elegans. PLOS ONE 10, e0124065 (2015).

34. Murareanu, B. M., Knox, J., Roy, P. J. & Reinke, A. W. High-throughput small molecule screen identifies inhibitors of microsporidia invasion and proliferation in *C. elegans*. bioRxiv 2021.09.06.459184 (2021) doi:10.1101/2021.09.06.459184.

35. Mok, C. A. et al. MIP-MAP: High-Throughput Mapping of Caenorhabditis elegans Temperature-Sensitive Mutants via Molecular Inversion Probes. Genetics 207, 447–463 (2017).

36. Almagro Armenteros, J. J., et al. SignalP 5.0 improves signal peptide predictions using deep neural networks. Nat. Biotechnol. 37, 420–423 (2019).

37. Reinke, A. W., Mak, R., Troemel, E. R. & Bennett, E. J. In vivo mapping of tissue- and subcellular-specific proteomes in Caenorhabditis elegans. Sci. Adv. 3, e1602426 (2017).

38. Ghafouri, S. & McGhee, J. D. Bacterial residence time in the intestine of Caenorhabditis elegans. Nematology 9, 87–91 (2007).

39. Engelmann, I. et al. A Comprehensive Analysis of Gene Expression Changes Provoked by Bacterial and Fungal Infection in C. elegans. PLoS ONE 6, e19055 (2011).

40. Head, B. P., Olaitan, A. O. & Aballay, A. Role of GATA transcription factor ELT-2 and p38 MAPK PMK-1 in recovery from acute *P. aeruginosa* infection in *C. elegans*. Virulence 8, 261–274 (2017).

41. Styer, K. L. et al. Innate immunity in Caenorhabditis elegans is regulated by neurons expressing NPR-1/GPCR. Science 322, 460–464 (2008).

42. Kim, D. H. et al. A Conserved p38 MAP Kinase Pathway in Caenorhabditis elegans Innate Immunity. Science 297, 623–626 (2002).

43. Tan, M. W., Mahajan-Miklos, S. & Ausubel, F. M. Killing of Caenorhabditis elegans by Pseudomonas aeruginosa used to model mammalian bacterial pathogenesis. Proc. Natl. Acad. Sci. U. S. A. 96, 715–720 (1999).

44. Kirienko, N. V., Cezairliyan, B. O., Ausubel, F. M. & Powell, J. R. Pseudomonas aeruginosa PA14 Pathogenesis in Caenorhabditis elegans. in Pseudomonas Methods and Protocols (eds. Filloux, A. & Ramos, J.-L.) 653–669 (Springer, 2014). doi:10.1007/978-1-4939-0473-0_50.

45. Cook, D. E., Zdraljevic, S., Roberts, J. P. & Andersen, E. C. CeNDR, the *Caenorhabditis elegans* natural diversity resource. Nucleic Acids Res. 45, D650–D657 (2017).

46. Dierking, K., Yang, W. & Schulenburg, H. Antimicrobial effectors in the nematode Caenorhabditis elegans: an outgroup to the Arthropoda. Philos. Trans. R. Soc. B Biol. Sci. 371, 20150299 (2016).

47. Suh, J. & Hutter, H. A survey of putative secreted and transmembrane proteins encoded in the C. elegans genome. BMC Genomics 13, 333 (2012).

48. Gallotta, I. et al. Extracellular proteostasis prevents aggregation during pathogenic attack. Nature 584, 410–414 (2020).

49. Strzyz, P. Bend it like glycocalyx. Nat. Rev. Mol. Cell Biol. 20, 388–388 (2019).

50. Hoffman, C. L., Lalsiamthara, J. & Aballay, A. Host Mucin Is Exploited by Pseudomonas aeruginosa To Provide Monosaccharides Required for a Successful Infection. mBio 11, (2020).

51. Steentoft, C. et al. Precision mapping of the human O-GalNAc glycoproteome through SimpleCell technology. EMBO J. 32, 1478–1488 (2013).

52. Jensen, P. H., Kolarich, D. & Packer, N. H. Mucin-type O-glycosylation – putting the pieces together. FEBS J. 277, 81–94 (2010).

53. Tran, D. T. & Ten Hagen, K. G. Mucin-type O-Glycosylation during Development. J. Biol. Chem. 288, 6921–6929 (2013).

54. Schulenburg, H. & Felix, M.-A. The Natural Biotic Environment of Caenorhabditis elegans. Genetics 206, 55–86 (2017).

55. Samuel, B. S., Rowedder, H., Braendle, C., Félix, M.-A. & Ruvkun, G. Caenorhabditis elegans responses to bacteria from its natural habitats. Proc. Natl. Acad. Sci. 113, E3941–E3949 (2016).

56. Sinha, A., Rae, R., Iatsenko, I. & Sommer, R. J. System Wide Analysis of the Evolution of Innate Immunity in the Nematode Model Species Caenorhabditis elegans and Pristionchus pacificus. PLOS ONE 7, e44255 (2012).

57. Ashe, A. et al. A deletion polymorphism in the Caenorhabditis elegans RIG-I homolog disables viral RNA dicing and antiviral immunity. eLife 2, e00994 (2013).

58. Reddy, K. C. et al. An Intracellular Pathogen Response Pathway Promotes Proteostasis in C. elegans. Curr. Biol. CB 27, 3544–3553.e5 (2017).

59. Toor, J. & Best, A. Evolution of Host Defense against Multiple Enemy Populations. Am. Nat. 187, 308–319 (2016).

60. Thaler, J. S., Fidantsef, A. L., Duffey, S. S. & Bostock, R. M. Trade-Offs in Plant Defense Against Pathogens and Herbivores: A Field Demonstration of Chemical Elicitors of Induced Resistance. J. Chem. Ecol. 25, 1597–1609 (1999).

61. Mok, C., Belmarez, G., Edgley, M. L., Moerman, D. G. & Waterston, R. H. PhenoMIP: High-Throughput Phenotyping of Diverse *Caenorhabditis elegans* Populations via Molecular Inversion Probes. G3 GenesGenomesGenetics 10, 3977–3990 (2020).

62. Sato, T. et al. The Rab8 GTPase regulates apical protein localization in intestinal cells. Nature 448, 366–369 (2007).

63. Troemel, E. R., Félix, M.-A., Whiteman, N. K., Barrière, A. & Ausubel, F. M. Microsporidia Are Natural Intracellular Parasites of the Nematode Caenorhabditis elegans. PLoS Biol. 6, e309 (2008).

64. Dunn, A. K., Millikan, D. S., Adin, D. M., Bose, J. L. & Stabb, E. V. New rfp- and pES213- derived tools for analyzing symbiotic Vibrio fischeri reveal patterns of infection and lux expression in situ. Appl. Environ. Microbiol. 72, 802–810 (2006).

65. Sifri, C. D., Begun, J., Ausubel, F. M. & Calderwood, S. B. Caenorhabditis elegans as a Model Host for Staphylococcus aureus Pathogenesis. Infect. Immun. 71, 2208–2217 (2003).

66. Moore, S. D. & Prevelige, P. E. A P22 scaffold protein mutation increases the robustness of head assembly in the presence of excess portal protein. J. Virol. 76, 10245–10255 (2002).

67. Ponchon, L., Beauvais, G., Nonin-Lecomte, S. & Dardel, F. A generic protocol for the expression and purification of recombinant RNA in Escherichia coli using a tRNA scaffold. Nat. Protoc. 4, 947–959 (2009).

68. Frøkjær-Jensen, C. et al. Single-copy insertion of transgenes in Caenorhabditis elegans. Nat. Genet. 40, 1375–1383 (2008).

69. Dickinson, D. J., Pani, A. M., Heppert, J. K., Higgins, C. D. & Goldstein, B. Streamlined Genome Engineering with a Self-Excising Drug Selection Cassette. Genetics 200, 1035– 1049 (2015).

70. Schindelin, J., et al. Fiji: an open-source platform for biological-image analysis. Nat. Methods 9, 676–682 (2012).

71. Thompson, O. et al. The million mutation project: a new approach to genetics in Caenorhabditis elegans. Genome Res. 23, 1749–1762 (2013).

72. Bolger, A. M., Lohse, M. & Usadel, B. Trimmomatic: a flexible trimmer for Illumina sequence data. Bioinforma. Oxf. Engl. 30, 2114–2120 (2014).

73. Li, H. & Durbin, R. Fast and accurate short read alignment with Burrows-Wheeler transform. Bioinforma. Oxf. Engl. 25, 1754–1760 (2009).

74. DePristo, M. A. et al. A framework for variation discovery and genotyping using next- generation DNA sequencing data. Nat. Genet. 43, 491–498 (2011).

75. Wang, K., Li, M. & Hakonarson, H. ANNOVAR: functional annotation of genetic variants from high-throughput sequencing data. Nucleic Acids Res. 38, e164 (2010).

76. Powell, J. R. & Ausubel, F. M. Models of Caenorhabditis elegans infection by bacterial and fungal pathogens. Methods Mol. Biol. Clifton NJ 415, 403–427 (2008).

77. Walhout, A. J. et al. GATEWAY recombinational cloning: application to the cloning of large numbers of open reading frames or ORFeomes. Methods Enzymol. 328, 575–592 (2000).

78. Hartley, J. L., Temple, G. F. & Brasch, M. A. DNA cloning using in vitro site-specific recombination. Genome Res. 10, 1788–1795 (2000).

79. Dokshin, G. A., Ghanta, K. S., Piscopo, K. M. & Mello, C. C. Robust Genome Editing with Short Single-Stranded and Long, Partially Single-Stranded DNA Donors in Caenorhabditis elegans. Genetics 210, 781–787 (2018).

80. Concordet, J.-P. & Haeussler, M. CRISPOR: intuitive guide selection for CRISPR/Cas9 genome editing experiments and screens. Nucleic Acids Res. 46, W242–W245 (2018).

81. Mello, C. C., Kramer, J. M., Stinchcomb, D. & Ambros, V. Efficient gene transfer in C.elegans: extrachromosomal maintenance and integration of transforming sequences. EMBO J. 10, 3959–3970 (1991).

82. Amrit, F. R. G., Ratnappan, R., Keith, S. A. & Ghazi, A. The C. elegans lifespan assay toolkit. Methods 68, 465–475 (2014).

83. Crittenden, S. & Kimble, J. Preparation and Immunolabeling of Caenorhabditis elegans. Cold Spring Harb. Protoc. 2009, pdb.prot5216-pdb.prot5216 (2009).

84. Wan, G. et al. Spatiotemporal regulation of liquid-like condensates in epigenetic inheritance. Nature 557, 679–683 (2018).

85. Large-Scale Screening for Targeted Knockouts in the Caenorhabditis elegans Genome. G3 GenesGenomesGenetics 2, 1415–1425 (2012).

